# Diminished signal-to-noise ratio disrupts somatosensory population encoding and drives tactile hyposensitivity in the *Fmr1*^-/y^ autism model

**DOI:** 10.1101/2024.08.08.607129

**Authors:** Ourania Semelidou, Théo Gauvrit, Célien Vandromme, Alexandre Cornier, Anna Saint-Jean, Yves Le Feuvre, Melanie Ginger, Andreas Frick

## Abstract

Touch is essential for interacting with the world, and atypical tactile experience is a core feature of autism that profoundly affects daily life. However, we do not know the neural mechanisms of low-level tactile perception and their alterations in autism. Using a translational perceptual task, we recapitulate the multifaceted tactile features of autistic individuals in the *Fmr1*^-/y^ mouse model of autism, showing tactile hyposensitivity, interindividual variability, and unreliable responses. We reveal that impaired detection decoding in *Fmr1*^-/y^-hyposensitive mice stems from diminished single-neuron signal-to-noise ratio in the primary somatosensory cortex that leads to weak population encoding of the tactile stimulus and its detection. This manifests as reduced stimulus-dependent neural recruitment, impaired response precision, and disrupted ensemble dynamics. Decreasing neuronal hyperexcitability strengthens sensory encoding and improves tactile perception. This work provides a translational framework for probing neuronal-perceptual changes in neurodevelopmental conditions, reveals inter-individual variability in preclinical models, and uncovers the neural basis of tactile hyposensitivity in autism.

## Introduction

Two-thirds of autistic individuals present alterations in tactile perception which, together with other sensory features, constitute one of the core diagnostic criteria of the condition^1,2^. Tactile perception is crucial for exploring and interacting with the world, and fine touch—the ability to detect and discriminate vibration, pressure, slip, and texture—plays a role in object exploration and social interactions^3^. Consequently, tactile alterations strongly impact daily life in autism while also contributing to the development of other core symptoms^4–6^. These tactile features are highly heterogeneous across individuals, a characteristic reflected in the DSM-5 as “hyper- or hypo-reactivity to sensory input”^7–11^.

Tactile alterations encompass both differences in fine touch (perceptual sensitivity) and touch-related neuronal responses^12^. Objective measures of perceptual sensitivity showed that the detection of vibrotactile stimuli delivered to the fingers depends on the stimulus frequency, with hyposensitivity to low-level stimulation^13,14^ and typical detection of higher-frequency stimuli^15^ in autism. Conversely, low-level vibrotactile stimuli delivered to the forearm and representing the social submodality of somatosensation revealed hypersensitivity in autistic individuals^16^.

Tactile information is relayed from the periphery to the primary somatosensory cortex, where it is processed and integrated to create haptic percepts^17^. In autistic individuals, tactile stimulation revealed diminished^18^ or increased^19^ early responses, decreased late responses^19^, reduced GABA levels^20,21^, and different representations of tactile stimuli^22^ within the somatosensory cortex. While these neuronal responses are rarely combined with perceptual measures, reduced GABA levels have been correlated with tactile hyposensitivity in autistic individuals^20^. Conversely, GABAergic interneuron dysfunction in the whisker-related primary somatosensory cortex in *Shank3B*^-/-^ mice was linked to perceptual hypersensitivity to whisker stimulation^23^. Nonetheless, the cortical mechanisms underlying tactile perceptual hyposensitivity in autism remain unclear.

Leveraging the similarity between the somatosensory systems of the mouse forepaw and human hand^24–26^, we developed a highly translational platform to study the neural mechanisms of fine touch, focusing on the detection of low-level, innocuous stimuli relevant to daily life. We show that stimulus encoding in the primary somatosensory cortex (S1-FP) mediates perception and is sufficient to decode behavioral responses. Through this standardized approach for rodent models, we further tackle the nuanced sensory symptomatology of autism and uncover the neural correlates of altered perceptual sensitivity. Using the *Fmr1*^-/y^ mouse model^27^, we recapitulate the multifaceted tactile features of autism, revealing high interindividual variability in tactile perception with a subgroup exhibiting tactile hyposensitivity and unreliable responses. In these hyposensitive *Fmr1*^-/y^ mice, S1-FP activity fails to reliably decode stimulus detection due to reduced single-neuron signal-to-noise ratio that leads to diminished population encoding of the tactile stimulus and its detection. Importantly, reducing neuronal hyperexcitability strengthens sensory encoding and improves tactile perception, uncovering a neural mechanism underlying tactile issues in autism.

## Results

### Forepaw-based perceptual decision-making task assesses tactile detection in mice

To study the perception of low-level vibrotactile stimuli, we reverse-translated a task used in human studies^13,14^ and developed a Go/No-Go detection task in head-fixed mice. We combined this task with two-photon calcium imaging to examine the neural correlates of tactile perception within the primary somatosensory neocortex (S1). Water-restricted *Fmr1*^-/y^ mice and their wild-type (WT) littermates were trained to associate the delivery of a fixed, suprathreshold vibrotactile stimulus (15 µm, 40 Hz) with a water reward (Go trial; Fig. 1a-left). During Go trials, vibrotactile stimuli (500 ms) were delivered to the right forepaw of the mice, followed by a 2-s response window during which the mice could lick to trigger water reward delivery (Fig. 1a-left). No-Go (catch) trials in which no stimulus was presented were used to assess spontaneous licking (Fig. 1a-right).

**Figure 1.**
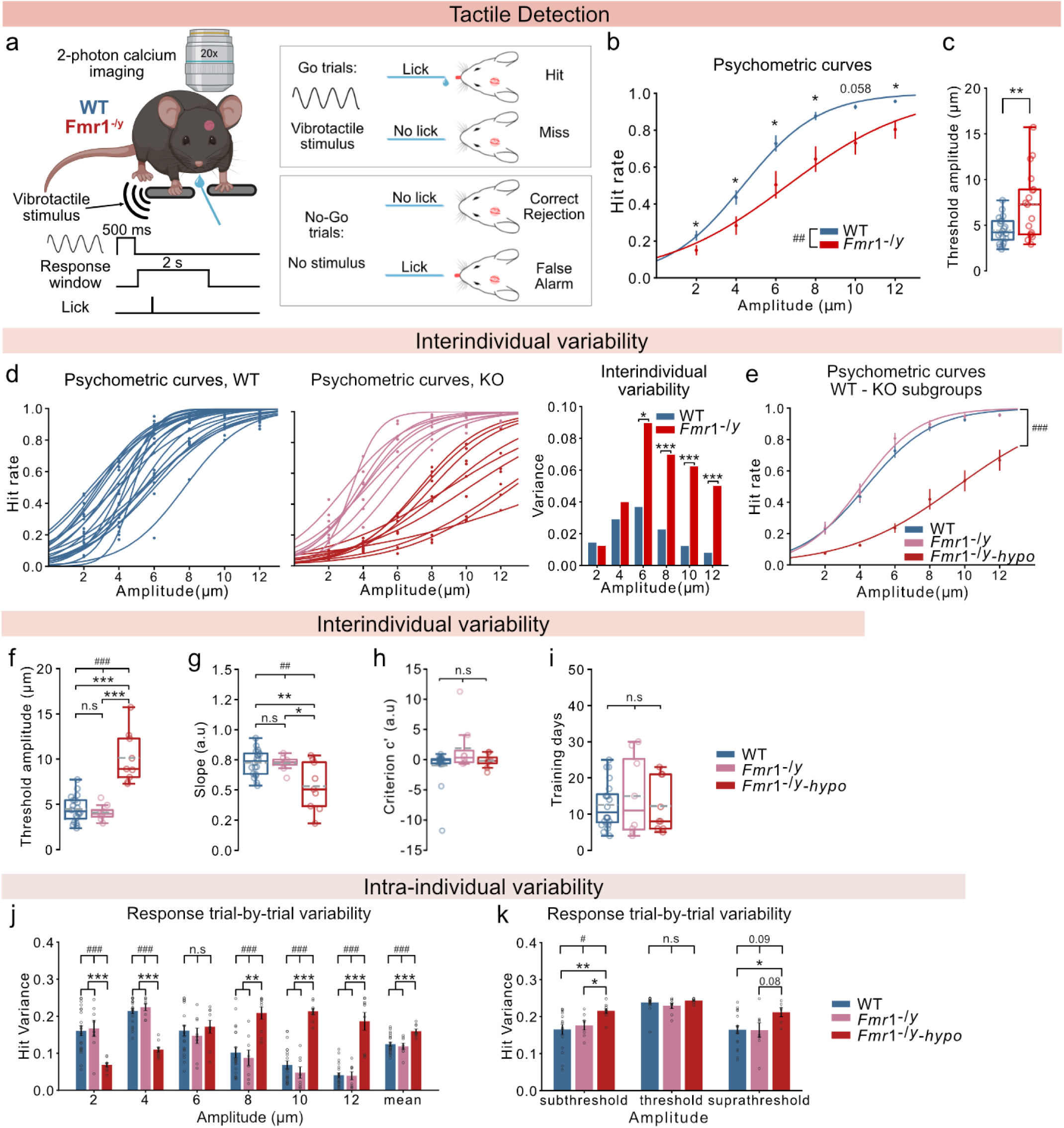
A forepaw-based perceptual decision-making task to assess tactile detection in mice. For panels **b, c, d, e, f, g, i, j,:** n=20 WT, 17 *Fmr1*^-/y^ mice (of which 9 *Fmr1*^-/y^-hyposensitive), 5-7 sessions with ∼130 repetitions of each amplitude per mouse. For panel **h,:** n=19 WT, 8 *Fmr1*^-/y^, 8 *Fmr1*^-/y^-hyposensitive mice. For panel **k,:** subthreshold: n=16 WT, 7 *Fmr1*^-/y^, 9 *Fmr1*^-/y^-hyposensitive mice, threshold: n=20 WT, 8 *Fmr1*^-/y^, 8 *Fmr1*^-/y^-hyposensitive mice, suprathreshold: n=20 WT, 8 *Fmr1*^-/y^,6 *Fmr1*^-/y^-hyposensitive mice. **a,** Left: schema showing the behavioral setup and trial protocol. Right: behavioral outcomes depending on the animal’s response. **b,** Psychometric curves fitted on average Hit rates for each stimulus amplitude. **c,** Detection thresholds calculated based on the psychometric curves. **d,** Psychometric curves of each individual mouse (left and middle, colors depict different groups) and variance of Hit rates (right). **e,** Psychometric curves fitted on average Hit rates for each stimulus amplitude for WT mice and the two Fmr1^-/y^ subgroups. Detection thresholds **(f,)** and perceptual accuracy **(g,)** calculated based on the psychometric curves in **(e,). h**, Relative criterion c’ depicting the licking strategy. **i,** Number of days mice spent in pre-training and training. **j,** Trial-by-trial variability calculated as response variance of Hit for each stimulus amplitude and the average for all amplitudes. **k,** Trial-by-trial variability of Hit responses following normalization for the threshold amplitude of each mouse. Subthreshold stimulus is the first stimulus with amplitude lower than the threshold and suprathreshold stimulus is the first amplitude higher than the threshold. P values were computed using Mixed ANOVA for panels **b, e,;** one-way ANOVA for panels **f, g, j, k,;** Kruskal-Wallis ANOVA for panels **h, i,;** two-sided t-test for panels **c, e, f, g, i, j, k,**; Mann-Whitney test for panels **b, e, h, i, j, k,**; F test for panel **d**,. ***/^###^P < 0.001, **/^##^P < 0.01, */^#^P < 0.05 or n.s, not significant. ^#^indicates results from ANOVA

To evaluate sensory perception, we quantified four behavioral outcomes: Hit for successful and Miss for unsuccessful Go trials, and Correct Rejection (CR) for successful and False Alarm (FA) for unsuccessful No-Go trials (Fig. 1a-right). Mice completed the training period when they met our learning criterion, with a performance exceeding an average 70% Hit rate with less than a 30% False Alarm rate for three consecutive training days (Fig. S1a). Pre-stimulus licking (3-8 s before stimulus onset) canceled stimulus delivery and led to a 5-s timeout. 86% of the WT mice and 73% of the *Fmr1*^-/y^ mice reached the learning criterion, with no difference between the learning performances of the two groups (Fig. S1b). Similarly, no genotype effect was observed for the training duration of mice that learned the task (Fig. S1c). These data show that both WT and *Fmr1*^-/y^ mice can learn the detection task, with similar success rates and training duration.

### *Fmr1*^-/y^ mice show perceptual hyposensitivity for low-level vibrotactile stimuli

Autistic individuals exhibit alterations in fine touch, with hyposensitivity to low-level (<25 µm amplitude, 25 Hz frequency) vibrotactile stimulation delivered to the fingers^13,14^. We tested whether we could replicate this phenotype in a preclinical model of autism using similar stimuli in our translational task. We assessed the tactile sensitivity of mice that met the learning criterion during the training period by delivering novel vibrotactile stimuli of low, fixed frequency (10 Hz) at 6 different amplitudes (range: 2-12 µm, 2 µm interval) in a pseudorandom manner. Mice reported the detection of a given stimulus by licking to receive a water reward. By analyzing the successful licks (Hit rates) for all stimulus amplitudes of sessions with low spontaneous licking (<40%), we generated psychometric curves (Fig. 1b) and determined the detection threshold (stimulus amplitude detected in 50% of the trials; curve inflection point), perceptual accuracy (curve slope), and licking strategy (criterion c’; i.e., how liberally or conservatively the mice licked while trying to get the reward) of each mouse.

*Fmr1*^-/y^ mice exhibited decreased tactile sensitivity (Hit rates) across the entire amplitude range (Fig. 1b), resulting in increased detection thresholds (Fig. 1c) and a trend for lower perceptual accuracy (Fig. S1d) compared to their WT littermates. This difference in detection was not due to alternative strategies adopted by the mice to obtain the reward, as indicated by the similar values of the criterion c’ (Fig. S1e).

Our results recapitulate the findings in autistic individuals, with *Fmr1*^-/y^ mice showing perceptual hyposensitivity to low-level vibrotactile forepaw stimulation.

### *Fmr1*^-/y^ mice show increased interindividual variability with a subgroup of mice exhibiting hyposensitivity during tactile detection

High interindividual variability in behavior and in neural responses to sensory stimuli is well-documented in autistic individuals^19,28–31^, but has not been recapitulated in mouse models of autism. We examined whether differences in interindividual variability can also be observed in our mouse model by comparing the variance of the Hit rates across amplitudes. *Fmr1*^-/y^ mice showed markedly increased interindividual variability compared to WT mice, for all stimuli above 4 µm (Fig. 1d).

Further inspection of the responses of *Fmr1*^-/y^ mice suggested the existence of discrete subgroups. We therefore used K-means clustering on the Hit rates across amplitudes to divide the *Fmr1*^-/y^ cohort into two subgroups. Consistent with findings from autistic individuals that revealed subgroups based on sensory difficulties^31^, one subgroup of *Fmr1*^-/y^ mice exhibited tactile stimulus detection comparable to that of WT mice, while the second subgroup displayed strongly diminished tactile sensitivity across all stimulus amplitudes (here referred to as *Fmr1*^-/y^-hyposensitive mice; Fig. 1e). *Fmr1*^-/y^-hyposensitive mice exhibited elevated perceptual detection thresholds (Fig. 1f) and decreased perceptual accuracy (cf. *Fmr1*^-y^ and WT group) (Fig. 1g). Importantly, no genotype differences were observed in strategy (Fig. 1h) and learning abilities (Fig. 1i, Fig. S1f).

Intrigued by the perceptual differences within the *Fmr1*^-/y^ cohort, we explored possible causes for this phenotypic heterogeneity. We first verified that mice of both subgroups came from the same litters, and controlled for the age of the parents and the experience of the mothers, thus excluding the possibility of litter- or breeder-specific variability^32^. No difference was observed in the litter size, a feature that correlates with behavioral alterations in mice^33^ (Fig. S1g). By constrast, *Fmr1*^-/y^-hyposensitive mice weighed on average 2.5 g more than the WT animals before water restriction, a difference that was not observed between WT mice and the other *Fmr1*^-/y^ subgroup (Fig. S1h). Body weight inversely correlates with sensitivity to visual stimuli in vertebrates^34^, as does the metabolic rate in humans^35^, and metabolic alterations have previously been described in the context of autism^36–38^.

These results demonstrate that interindividual variability can be recapitulated in mouse models of autism and reveal a subgroup in the *Fmr1*^-/y^ cohort with tactile hyposensitivity.

### *Fmr1*^-/y^-hyposensitive mice exhibit increased intra-individual variability across stimulus trials

Increased intra-individual variability has been observed in the behavioral and neural responses of autistic individuals^8,39–43^. However, this feature has not been examined in the context of sensory detection in autism. Our results on perceptual accuracy among *Fmr1*^-/y^-hyposensitive mice (Fig. 1g) suggest less reliable sensory perception. To further investigate this finding, we assessed the trial-by-trial (TBT) variability in the behavioral response for each stimulus amplitude in Go trials (Hit vs Miss). As expected for WT mice and *Fmr1*^-/y^ mice with intact perception, the response variability increased close to threshold stimulation (Fig. 1j). Conversely, *Fmr1*^-/y^-hyposensitive mice exhibited increased TBT variability for all stimulus amplitudes above 6 µm, and increased mean variance of responses across all amplitudes compared to the other groups (Fig. 1j).

Since the threshold amplitude differs between mice and intra-individual variability is elevated at the threshold, we examined genotype differences in response variability specifically for threshold and non-threshold stimuli. We compared the TBT variability of responses at the threshold amplitude (50% Hit rate) and the first sub-threshold and supra-threshold amplitudes for each mouse. As expected, all groups of mice showed a large TBT variability in their responses to threshold stimuli. However, *Fmr1*^-/y^-hyposensitive mice exhibited significantly more variable responses for sub- and suprathreshold stimulus amplitudes (Fig. 1k). These data indicate that unreliable responses in autism extend to tactile detection, showing high intra-individual variability in *Fmr1*^-/y^-hyposensitive mice across a spectrum of amplitudes, from sub- to supra-threshold levels.

### Neuronal activity in the S1-FP fails to predict stimulus detection in *Fmr1*^-/y^-hyposensitive mice

Are low-level stimuli encoded differently in the cortex of *Fmr1*^-/y^-hyposensitive mice, leading to impaired tactile detection? To address this question, we explored the neural correlates of low-level vibrotactile detection in the forepaw-related primary somatosensory cortex (S1-FP) and their potential changes in autism. We recorded single-cell activity from populations of excitatory pyramidal neurons and inhibitory GABAergic interneurons in S1-FP of mice engaged in the tactile detection task (Fig. 2a). Neuronal activity was aligned to tactile stimulation and behavioral response (licking or absence thereof), allowing us to dissect how sensory stimulation and its detection are represented in local cortical circuits (Fig. 2b). The total number of recorded pyramidal neurons and inhibitory interneurons in the field of view was in the range of 50-150 (∼20% interneurons, no genotype difference, Fig. S2a-b).

**Figure 2.**
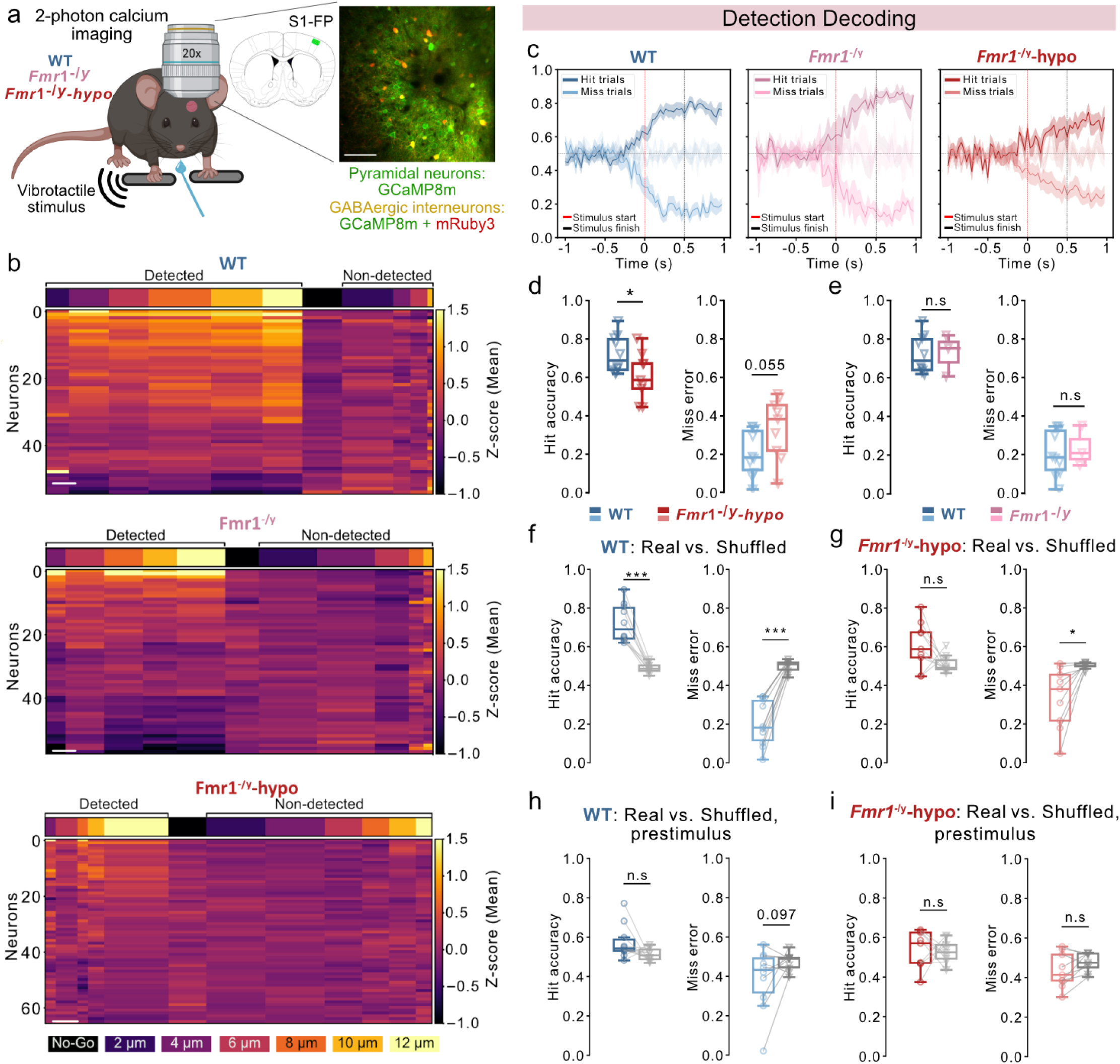
Detection decoding in S1-FP. For panels **c-i**: n=10 WT, 3 *Fmr1*^-/y^ mice with intact tactile thresholds, 9 *Fmr1*^-/y^-hyposensitive mice.1 session, ∼10 repetitions of each amplitude per mouse. **a,** Left: schema showing the imaging setup during tactile detection. Right: Position of injections in the forepaw-related primary somatosensory cortex and example of the field of view (scale bar: 100 mm). **b,** Example heat maps of activity (z-score) during stimulus delivery for one WT, one *Fmr1*^-/y^, and one *Fmr1*^-/y^-hyposensitive mouse. Each horizontal line depicts a neuron. Vertical lines show averaged activity across trials for each stimulus amplitude. Stimuli are sorted depending on their amplitude and whether they were detected (left) or not (right). No-Go trials are shown in the middle (black). Color-coding of each amplitude is shown at the bottom. (scale bar: 5 trials) **c,** Response decoding in each mouse group. Stimulus start is indicated by a red dashed line and stimulus end by a black dashed line. The dark line represents true Hit classification and the light line represents false Miss (Miss error) classification. Datasets including prestimulus activity (1 s), tactile stimulation (500 ms), post-stimulus activity (500 ms) were used to train the logistic regression model. Classifiers were trained on each frame to decode if the trial was detected (Hit) or non-detected (Miss) (adapted from ref^44^). Comparison of the average Hit accuracy (left) and Miss error (right) of the classifiers during stimulation between WT and *Fmr1*^-/y^-hyposensitive mice **(d,)** and between WT and *Fmr1*^-/y^ mice with intact detection thresholds **(e,).** Average Hit accuracy (left) and Miss error (right) of the classifiers on the total duration of the stimulation (500 ms) compared to the accuracy of the model trained with shuffled behavioral labels in WT mice **(f,)** and *Fmr1*^-/y^-hyposensitive mice **(g,)**. Average Hit accuracy (left) and Miss error (right) of the classifiers on the prestimulus activity (200 ms) compared to the accuracy of the model trained with shuffled behavioral labels in WT mice **(h,)** and *Fmr1*^-/y^-hyposensitive mice **(i,)**. P values were computed using two-sided t-test for panel **d, e,;** two-sided paired t-test for panel **f, g, h-right, I,;** Wilcoxon signed-rank test for panels **h-left,.** ***P<0.001, *P < 0.05 or n.s, not significant.

We first asked whether stimulus detection could be predicted from S1-FP population activity. To decode the transformation of sensory input into perceptual decisions, we implemented a frame-by-frame logistic regression classifier^44^ trained on neuronal activity during both the pre-stimulus (1 s) and stimulus (500 ms) windows (32 ms time bins; Fig. 2c). Our model successfully distinguished detected from non-detected trials in WT mice and in *Fmr1*^-/y^ mice with typical detection thresholds, but performed worse in *Fmr1*^-/y^-hyposensitive mice (Fig. 2c).

Direct genotype comparisons of decoding accuracy during stimulus presentation revealed a significant reduction in the model’s accuracy to predict detected trials (Fig. 2d-left) and a strong trend toward increased errors for non-detected trials in *Fmr1*^-/y^-hyposensitive mice (Fig. 2d-right). In contrast, decoding performance was comparable between WT mice and *Fmr1*^-/y^ mice with intact detection thresholds, confirming that impaired decoding is specifically associated with the hyposensitive phenotype (Fig. 2e). We therefore focused our subsequent analyses on WT and *Fmr1*^-/y^-hyposensitive animals.

To ensure that classifier performance reflected genuine stimulus-related coding rather than chance correlations, we compared the decoding accuracy against classifiers trained on data with shuffled behavioral labels. In WT mice, decoding accuracy was significantly higher than the shuffled control (Fig. 2f), confirming that tactile detection can be reliably decoded from S1-FP activity. By contrast, the classifiers failed to predict detection in *Fmr1*^-/y^-hyposensitive mice, with Hit accuracy comparable to the shuffled control (Fig. 2g). As expected, decoding failed in both genotypes when the model was trained on pre-stimulus activity (200 ms window; Fig. 2h–i), indicating that accurate decoding was specific to the stimulus-delivery period.

Altogether, our findings demonstrate that neuronal activity in S1-FP robustly encodes tactile detection in WT mice and *Fmr1*^-/y^ mice with intact detection thresholds, but provides a less reliable readout of stimulus detection in *Fmr1*^-/y^-hyposensitive mice, indicating a disrupted cortical representation.

### S1-FP pyramidal neurons dominate the decoding of stimulus detection

To dissect the contributions of distinct neuronal populations to detection decoding, we pooled datasets from both genotypes and trained logistic regression classifiers on the average z-scored activity of individual neurons during stimulus delivery. We compared model performance when trained on either all neurons, only pyramidal neurons, or only GABAergic interneurons. Decoding accuracy was comparable when using all neurons or only pyramidal cells (Fig. S3a), whereas models trained solely on inhibitory interneurons performed significantly worse (Fig. S3b-c), demonstrating that pyramidal neurons carry the majority of information relevant to stimulus detection. Notably, decoding accuracy was reduced in *Fmr1*^-/y^-hyposensitive mice regardless of cell type (Fig. S3d-e).

These findings demonstrate that stimulus detection in S1-FP is primarily encoded by pyramidal neurons, and that this process is disrupted in *Fmr1*^-/y^-hyposensitive mice.

### *Fmr1*^-/y^-hyposensitive mice show weak detection encoding in pyramidal neuron population

To determine whether reduced detection decoding stems from altered detection encoding at the single-neuron level, we computed the sensitivity index (d’) of individual neurons, quantifying their ability to discriminate between detected and non-detected trials. Both pyramidal neurons (Fig. S4a, top) and GABAergic interneurons (Fig. S4a, bottom) showed low detection sensitivity across genotypes, with mean d′ values below 0.25 (Fig. S4b) and no differences in the proportion of detection-encoding neurons (d’>1; 1-5% on average) (Fig. S4c). These findings demonstrate that stimulus detection is not represented at the single-neuron level, suggesting that it rather emerges from coordinated population activity.

To test this hypothesis, we assessed encoding at the population level by quantifying the proportion of pyramidal neurons and GABAergic interneurons that were recruited (i.e., activated or inhibited) during stimulus presentation (Fig. S4d) and detection (Fig. S4e). Across all groups, higher stimulus amplitudes recruited more pyramidal neurons and GABAergic interneurons. However, a significant group difference was found in pyramidal neuron recruitment during stimulus encoding (Fig. S4d-top) and a trend was observed during detection encoding (Fig. S4e-top), indicating altered pyramidal population encoding in S1-FP in *Fmr1*^-/y^-hyposensitive mice.

To directly assess detection encoding at the population level, we calculated the proportion of neurons recruited during detected versus non-detected trials across all stimulus amplitudes. Pyramidal neurons, which contribute more strongly to stimulus detection than GABAergic interneurons (Fig. S3a-c), showed a significantly higher recruitment during detected than non-detected trials across genotypes (Fig. 3a-b), confirming that perceptual detection is reflected in population-level pyramidal activity within S1-FP. However, this detection-specific enhanced recruitment was less pronounced in *Fmr1*^-/y^-hyposensitive mice (Fig. 3c), blunting stimulus detection. While GABAergic interneurons also showed higher recruitment during detected versus non-detected trials across genotypes (Fig. 3d-e), there was no genotype difference in this feature (Fig. 3f).

**Figure 3.**
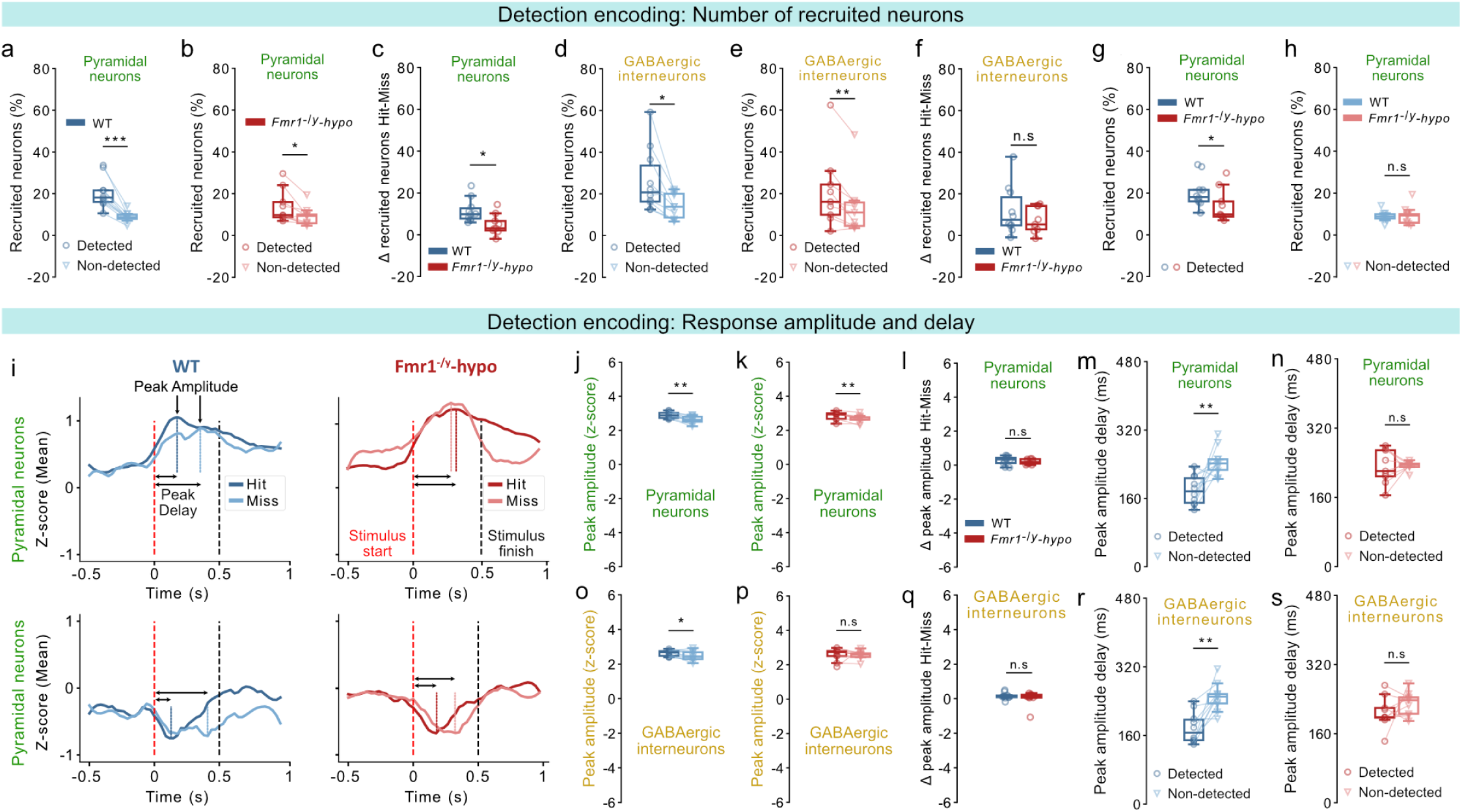
Detection encoding in S1-FP. For panels: n=10 WT, 9 *Fmr1*^-/y^-hyposensitive mice.1 session, ∼10 repetitions of each amplitude per mouse. Proportion of recruited (activated or inhibited) pyramidal neurons during detected and non-detected stimuli in WT mice **(a,)** and *Fmr1*^-/y^-hyposensitive mice **(b,)**. **c,** Difference in the proportion of recruited pyramidal neurons between detected and non-detected stimuli. **d,** Same as **a,** but for GABAergic interneurons. **e,** Same as **b,** but for GABAergic interneurons. **f,** Same as **c,** but for GABAergic interneurons. **g,** Proportion of recruited pyramidal neurons during detected stimuli. **h,** Same as **g,** but during non-detected stimuli. **i,** Calcium traces of pyramidal responses, averaged across trials for an example WT and *Fmr1*^-/y^-hyposensitive mouse. Z-score responses of pyramidal neurons before, during, and after stimulation during detected (Hit) and non-detected (Miss) trials are shown. Peak amplitude and peak delay are indicated. Quantification of peak response amplitude (z-score) of recruited pyramidal neurons during detected and non-detected stimuli in WT mice **(j,)** and *Fmr1*^-/y^-hyposensitive mice **(k,)**. **l,** Difference in the peak amplitude (z-score) of recruited pyramidal neurons between detected and non-detected stimuli. Quantification of response delay (ms) of the peak amplitude relative to stimulus onset for recruited pyramidal neurons during detected and non-detected stimuli in WT mice **(m,)** and *Fmr1*^-/y^-hyposensitive mice **(n,)**. **o,** Same as **j,** but for GABAergic interneurons. **p,** Same as **k,** but for GABAergic interneurons. **q,** Same as **l,** but for GABAergic interneurons. **r,** Same as **m,** but for GABAergic interneurons. **s,** Same as **n,** but for GABAergic interneurons. P values were computed using two-sided t-test for panels **c, f, h, l,**; two-sided paired t-test for panel **a, d, j, k, m, o, p, r, s,**; Wilcoxon signed-rank test for panels **b, e, n,**; Mann-Whitney U Test for panels **g, q,**. ***P<0.001, **P < 0.01, *P < 0.05 or n.s, not significant.

During detected trials, *Fmr1*^-/y^-hyposensitive mice showed significantly reduced recruitment of pyramidal neurons (Fig. 3g), whereas no differences were observed during non-detected trials (Fig. 3h). These differences could not be attributed to licking intention, as recruitment during False Alarm No-Go trials was comparable across genotypes (Fig. S4f-g). In line with our previous results (Fig. 3f), the recruitment of GABAergic interneurons did not differ between genotypes for either detected (Fig. S4h) or non-detected trials (Fig. S4i).

Together, these findings demonstrate that reduced decoding performance in *Fmr1*^-/y^-hyposensitive mice stems from weak detection encoding in pyramidal neuron populations, where reduced recruitment leads to less distinct responses between detected and non-detected trials.

### Reduced temporal sharpening contributes to weak detection encoding in *Fmr1*^-/y^-hyposensitive mice

Beyond the number of recruited neurons, the strength and timing of neuronal responses may further enhance stimulus detection encoding^45^. To explore this, we compared the peak response amplitude of recruited neurons and its delay relative to stimulus onset during detected and non-detected trials (Fig. 3i). In pyramidal neurons, response amplitudes were significantly higher during detected trials across genotypes (Fig. 3j-k), with no genotype differences in gain modulation (Fig. 3l). Whereas WT mice also exhibited faster pyramidal responses during detected trials (Fig. 3m), this temporal sharpening was absent in *Fmr1*^-/y^-hyposensitive mice (Fig. 3n).

As with pyramidal neurons, GABAergic interneurons in WT mice showed larger response amplitudes in detected trials (Fig. 3o), while *Fmr1*^-/y^-hyposensitive mice displayed no amplitude difference across trial types (Fig. 3p). However, gain modulation magnitude was comparable across genotypes (Fig. 3q), suggesting only a small deficit in *Fmr1*^-/y^-hyposensitive mice. Consistent with pyramidal neurons, response latencies in *Fmr1*^-/y^-hyposensitive mice were similar across trial types, indicating reduced inhibitory temporal sharpening (Fig. 3r-s).

These findings demonstrate that detection encoding is reinforced by both amplified response strength and faster response dynamics. Reduced temporal sharpening in *Fmr1*^-/y^-hyposensitive mice contributes to their impaired cortical representation of detected stimuli.

### Disrupted network reliability weakens ensemble encoding of tactile stimulation in *Fmr1*^-/y^-hyposensitive mice

Coordinated activity within neuronal ensembles is critical for efficient sensory perception^46–49^. To assess the relevance of this feature during the processing of low-level tactile information within S1-FP, we analyzed pairwise correlations in the recruitment of neurons during stimulus delivery (Fig. 4a). *Fmr1*^-/y^-hyposensitive mice exhibited significantly reduced response correlations among the pyramidal neuron (Fig. 4b) but not GABAergic interneuron ensembles (Fig. 4c). Neural correlations during No-Go catch trials were similar across genotypes for both cell types (Fig. S4j-k). These results show that disrupted pyramidal ensemble dynamics during tactile stimulation in *Fmr1*^-/y^-hyposensitive mice impede stimulus encoding.

**Figure 4.**
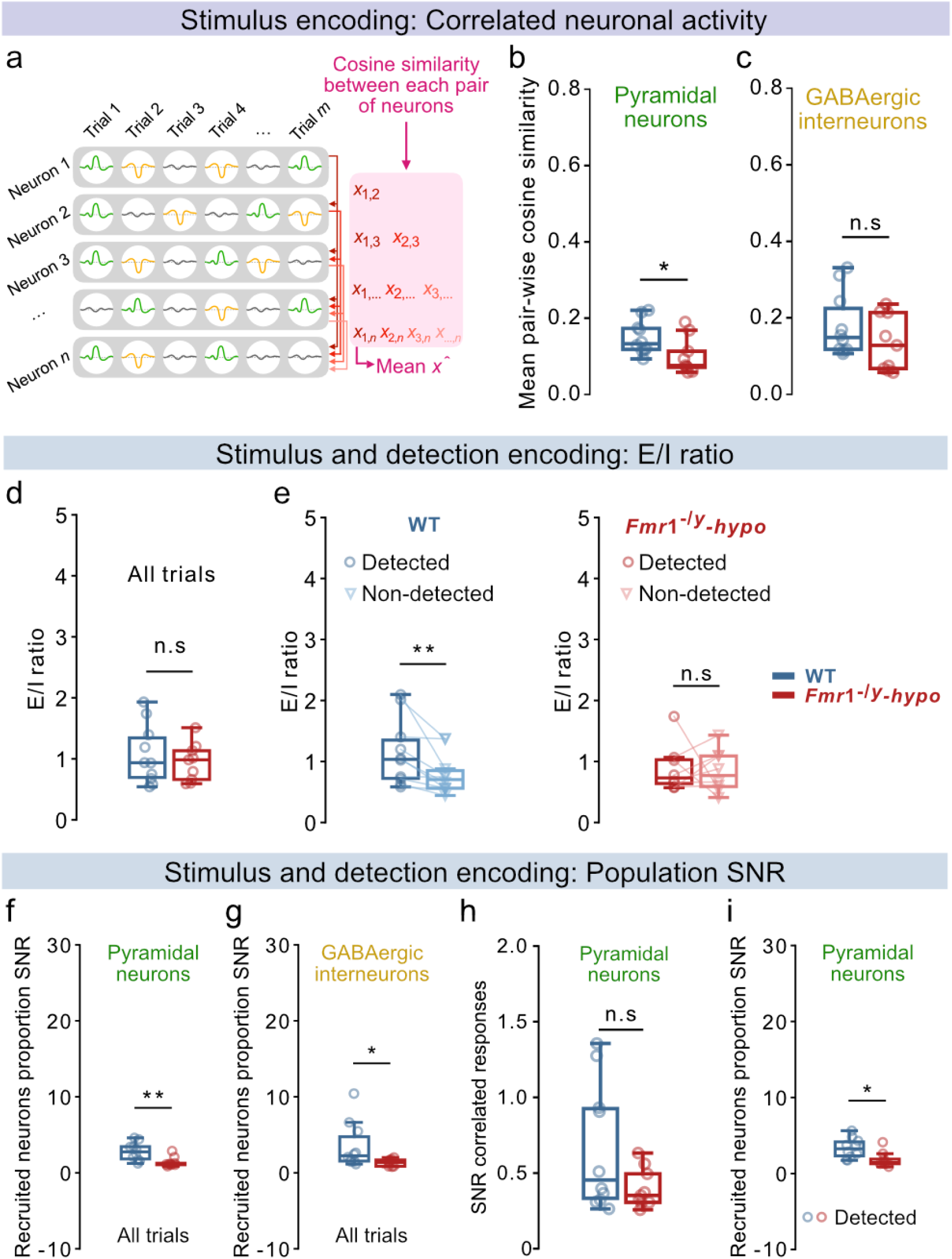
Neural response correlation and population signal-to-noise ratio (SNR) during stimulus and detection encoding. For panel **e,**: n=10 WT, 8 *Fmr1*^-/y^-hyposensitive mice, for all other panels: n=10 WT, 9 *Fmr1*^-/y^-hyposensitive mice.1 session, ∼10 repetitions of each amplitude per mouse **a,** Schema illustrating the calculation of mean pairwise cosine similarity, used to quantify trial-to-trial correlated activity between neuron pairs. **b,** Mean pair-wise cosine similarity of pyramidal trial-by-trial responses during stimulation. **c,** Same as **b,** but for GABAergic interneurons. **d,** Ratio of the proportion of activated pyramidal neurons to that of activated GABAergic interneurons during stimulus presentation. **e,** Ratio of the proportion of activated pyramidal neurons to that of activated GABAergic interneurons during detected and non-detected trials. **f,** Ratio of the proportion of recruited pyramidal neurons during stimulation to that during No-Go (catch) trials. **g,** Same as **f,** but for GABAergic interneurons. **h,** Ratio of the mean pair-wise correlation among pyramidal neurons during tactile stimulation to that during No-Go (catch) trials. **i,** Ratio of the proportion of recruited pyramidal neurons during detected Go trials to that during No-Go (catch) trials. P values were computed using two-sided t-test for panels **d,**; Wilcoxon signed-rank test for panel **e,**; Mann-Whitney U Test for panels **b, c, f, g, h, i,**. **P < 0.01, *P < 0.05 or n.s, not significant.

Diminished neural synchrony in *Fmr1*^-/y^-hyposensitive mice may reflect an altered excitatory-inhibitory ratio^50^. We tested this hypothesis by quantifying the ratio of activated pyramidal neurons to that of activated GABAergic interneurons during stimulus delivery. While E/I ratios across all trials did not differ between genotypes (Fig. 4d), WT mice displayed a clear increase in E/I ratio during detected trials compared to non-detected trials (Fig. 4e-left). This detection-dependent modulation of E/I balance was absent in *Fmr1*^-/y^-hyposensitive mice, where E/I ratios remained similar regardless of perceptual outcome (Fig. 4e-right).

Together, these findings demonstrate that disrupted pyramidal ensemble activity in S1-FP contributes to the impaired cortical encoding of tactile stimuli in *Fmr1*^-/y^-hyposensitive mice and that reduced detection encoding is associated with a lack of detection-dependent E/I modulation.

### Reduced population signal-to-noise ratio is linked to impaired stimulus and detection encoding in *Fmr1*^-/y^-hyposensitive mice

Effective sensory encoding relies on a high signal-to-noise ratio (SNR), which reflects the magnitude of the stimulus-evoked activity relative to ongoing background activity. To assess this, we quantified the SNR at the population level by comparing the proportion of recruited neurons during stimulation in Go trials versus the proportion of neurons that were spontaneously activated or inhibited during No-Go (catch) trials. *Fmr1*^-/y^-hyposensitive mice showed a markedly smaller increase in population SNR during stimulation for both pyramidal neurons (Fig. 4f) and GABAergic interneurons (Fig. 4g). In contrast, no genotype differences were observed in the ratio of correlated ensemble responses between stimulus delivery and No-Go trials for either pyramidal neurons (Fig. 4h) or GABAergic interneurons (Fig. S5a). These results demonstrate that stimulus-evoked responses at the population level were less distinct from ongoing neural recruitment in *Fmr1*^-/y^-hyposensitive mice.

Is this reduced SNR also linked to impaired detection encoding in *Fmr1*^-/y^-hyposensitive mice? Both groups exhibited increased population SNR during detected trials compared to non-detected ones, for both cell types (Fig. S5b-c). However, during detected trials, population SNR was significantly smaller in pyramidal neurons in *Fmr1*^-/y^-hyposensitive compared to WT mice (Fig. 4i), with no genotype differences in GABAergic interneuron SNR (Fig. S5d). In non-detected trials, population SNR was comparable across genotypes for pyramidal neurons (Fig. S5e) but reduced for GABAergic interneurons in *Fmr1*^-/y^-hyposensitive mice (Fig. S5f), indicating decreased subthreshold sensory gating due to reduced feedforward inhibition^20^. These results show that reduced pyramidal population SNR impacts detection encoding in *Fmr1*^-/y^-hyposensitive mice.

Together, these findings reveal a pervasive reduction in pyramidal SNR at the population level in *Fmr1*^-/y^-hyposensitive mice, disrupting reliable cortical encoding of the tactile stimulus and its detection.

### Reduced single-neuron signal-to-noise ratio underlies diminished neural recruitment in *Fmr1*^-/y^-hyposensitive mice

Low *population* SNR in terms of neural recruitment during stimulation suggests that reduced *single-neuron* SNR in *Fmr1*^-/y^-hyposensitive mice impairs the recruitment of sufficient pyramidal neuron numbers required for efficient encoding of tactile stimuli and their detection. To test this hypothesis, we calculated single-neuron SNR as the difference between the z-scored neural responses during stimulus delivery and those during No-Go catch trials for each neuron. During both stimulus and detection encoding, pyramidal neurons in *Fmr1*^-/y^-hyposensitive mice exhibited significantly lower single-neuron SNR compared to their WT littermates (Fig. 5a-b). In contrast, no differences were observed for GABAergic interneurons (Fig. 5c-d). Comparison of single-neuron SNR between detected and non-detected trials revealed an increased single-neuron SNR during detection across genotypes (Fig. S5i-j), mirroring our population SNR results (Fig. S5e-f).

**Figure 5.**
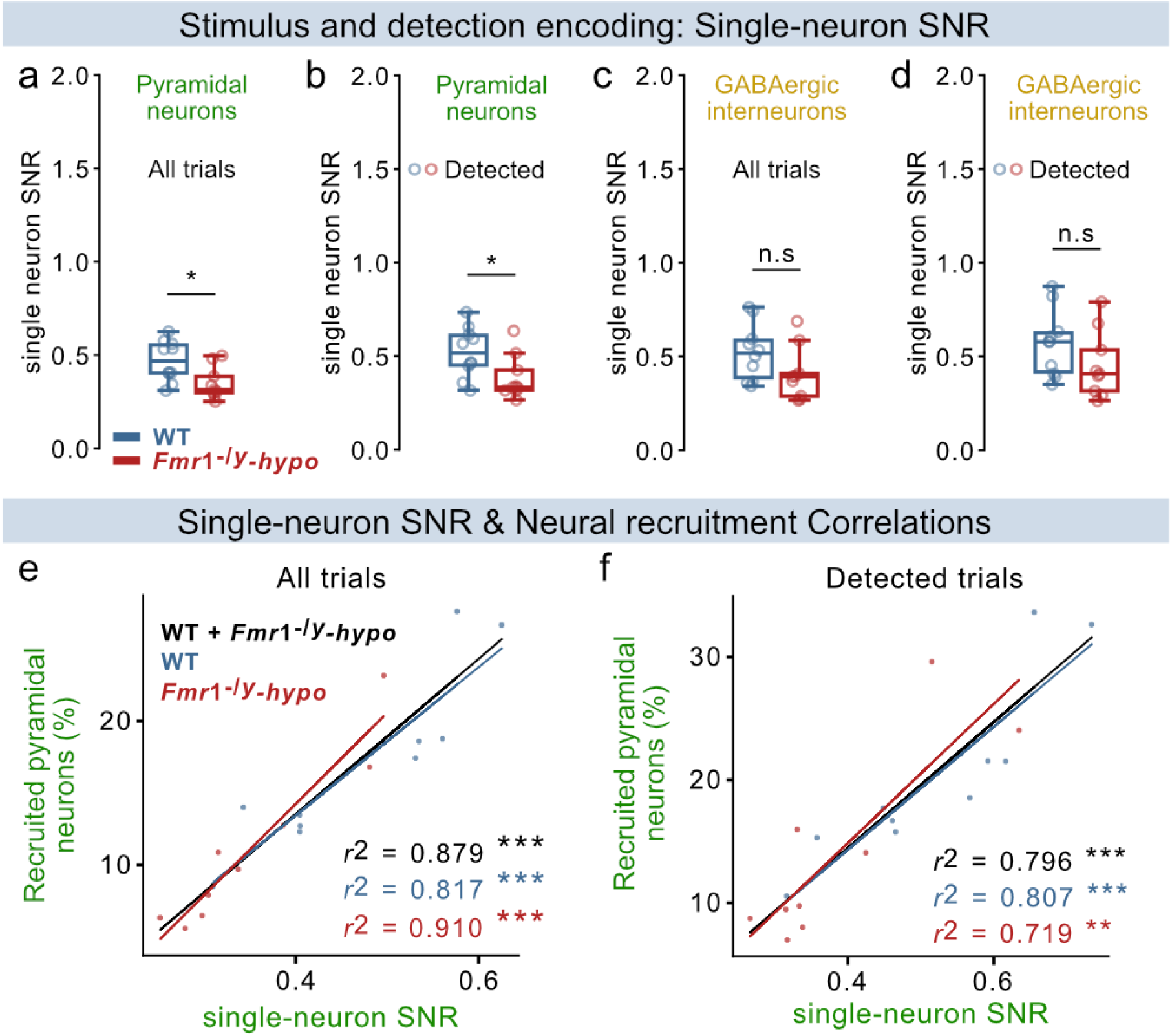
Reduced single-cell signal-to-noise ratio (SNR) during stimulus and detection encoding underlies diminished neural recruitment in *Fmr1*^-/y^-hyposensitive mice. For all panels: n=10 WT, 9 *Fmr1*^-/y^-hyposensitive mice. **a,** Difference between the average z-score response of pyramidal neurons during stimulation and during No-Go (catch) trials. **b,** Difference between the average z-score response of pyramidal neurons during detected stimuli and during No-Go (catch) trials. **c,** Same as **a,** but for GABAergic interneurons. **d,** Same as **b,** but for GABAergic interneurons. **e,** Correlation between single-neuron SNR and the proportion of stimulus-recruited pyramidal neurons during stimulation, computed for each animal. **f,** Correlation between single-neuron SNR and the proportion of stimulus-recruited pyramidal neurons during detected stimuli, computed for each animal. P values were computed using two-sided t-test for panels **a, c, d,**; Mann-Whitney U Test for panel **b,**; Linear least-squares regression for panels **e, f,**. ***P < 0.001, **P < 0.01, *P < 0.05 or n.s, not significant.

To test whether this decreased single-neuron SNR is indeed responsible for the reduced pyramidal recruitment observed in *Fmr1*^-/y^-hyposensitive mice during tactile stimulation, we correlated single-neuron SNR with the proportion of recruited neurons per animal. Pyramidal single-neuron SNR strongly correlated with recruitment during both stimulus (Fig. 5e) and detection encoding (Fig. 5f) across genotypes.

Together, these findings reveal that a strong reduction in single-cell pyramidal SNR underlies reduced recruitment of these neurons in *Fmr1*^-/y^-hyposensitive mice, disrupting reliable cortical encoding of the tactile stimulus and its detection.

### Decreased neuronal recruitment, signal-to-noise ratio, and ensemble dynamics in S1-FP lead to elevated detection thresholds in *Fmr1*^-/y^-hyposensitive mice

Behavioral analyses revealed significantly elevated detection thresholds in *Fmr1*^-/y^-hyposensitive mice (Fig. 1d, e-f). To investigate how this behavioral deficit is reflected in cortical processing alterations, we compared neuronal recruitment in S1-FP at the average detection threshold of WT mice (4 µm stimulus amplitude; Fig. 6a). At this stimulus amplitude, *Fmr1*^-/y^-hyposensitive mice exhibited reduced recruitment of pyramidal neurons (Fig. 6b), while their response amplitude (Fig. 6c) and timing (Fig. 6d) remained unchanged. Neuronal ensemble activity during stimulus encoding was also less correlated among pyramidal neurons (Fig. 6e), alongside reductions in population and single-neuron SNR (Fig. 6f-g). Comparable impairments were observed in GABAergic interneuron responses at the average WT detection threshold (Fig. 6h-m).

**Figure 6.**
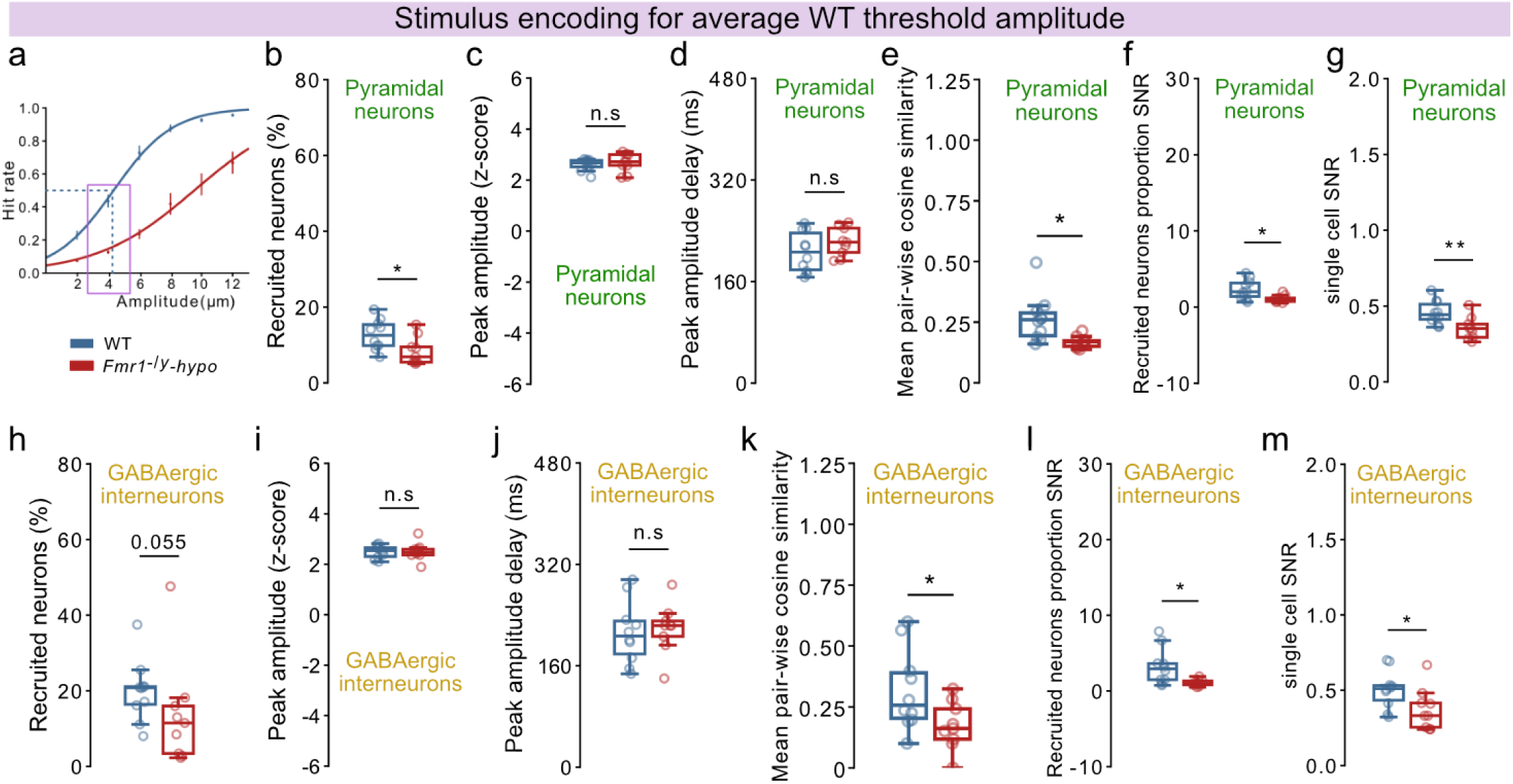
Reduced neural recruitment, SNR, and correlated responses in *Fmr1*^-/y^-hyposensitive mice result in increased detection thresholds. For all panels: n=10 WT, 9 *Fmr1*^-/y^-hyposensitive mice; analysis of responses during trials of 4 µm amplitude (average WT threshold). **a,** Schema showing the average WT threshold at 4 µm amplitude. **b,** Proportion of recruited pyramidal neurons. Peak amplitude (z-score) **(c,)** and peak amplitude delay **(d,)** of pyramidal responses. **e,** Mean pair-wise cosine similarity of pyramidal trial-by-trial responses. **f,** Ratio of the proportion of recruited pyramidal neurons during trials of 4 µm amplitude and during No-Go (catch) trials. **g,** Difference between the average z-score response of pyramidal neurons during trials of 4 µm amplitude and during No-Go (catch) trials. **h,** Same as **b,** but for GABAergic interneurons. **i,** Same as **c,** but for GABAergic interneurons. **j,** Same as **d,** but for GABAergic interneurons. **k,** Same as **e,** but for GABAergic interneurons. **l,** Same as **f,** but for GABAergic interneurons. **m,** Same as **g,** but for GABAergic interneurons. P values were computed two-sided t-test for panels **b, c, d, e, f, g, i, j, k, l, m,**; Mann-Whitney U Test for panel **h,**. **P < 0.01, *P < 0.05 or n.s, not significant.

Together, these findings directly link reduced single-neuron SNR and disrupted population-level evoked responses to behavioral hyposensitivity. Elevated detection thresholds in *Fmr1*^-/y^-hyposensitive mice arise from weakened cortical encoding of WT threshold stimuli, characterized by degraded SNR, reduced neuronal recruitment, and impaired ensemble coordination in both pyramidal and GABAergic neurons in S1-FP.

### Neuronal hyperexcitability contributes to tactile hyposensitivity in *Fmr1*^-/y^ mice

To gain insight into the cellular mechanisms underlying weak stimulus and detection encoding within the S1-FP and their contribution to atypical perception in *Fmr1*^-/y^-hyposensitive mice, we manipulated neuronal excitability by targeting the large conductance voltage- and calcium-sensitive potassium (BK_Ca_) channels. Dysfunction of BK_Ca_ channels has been associated with neurodevelopmental conditions such as autism and fragile X syndrome^51–53^, and administration of BK_Ca_ channel agonists can improve certain neuronal and behavioral alterations in *Fmr1*^-/y^ mice^54–57^. To test whether neuronal hyperexcitability contributes to tactile perception alterations of *Fmr1*^-/y^-hyposensitive mice, we administered the brain blood barrier-permeable BK_Ca_ channel agonist, BMS-204352 (BMS), or its vehicle, via intraperitoneal injection 30-45 minutes before tactile detection testing (Fig. 7a), following established protocols^54,55^.

**Figure 7.**
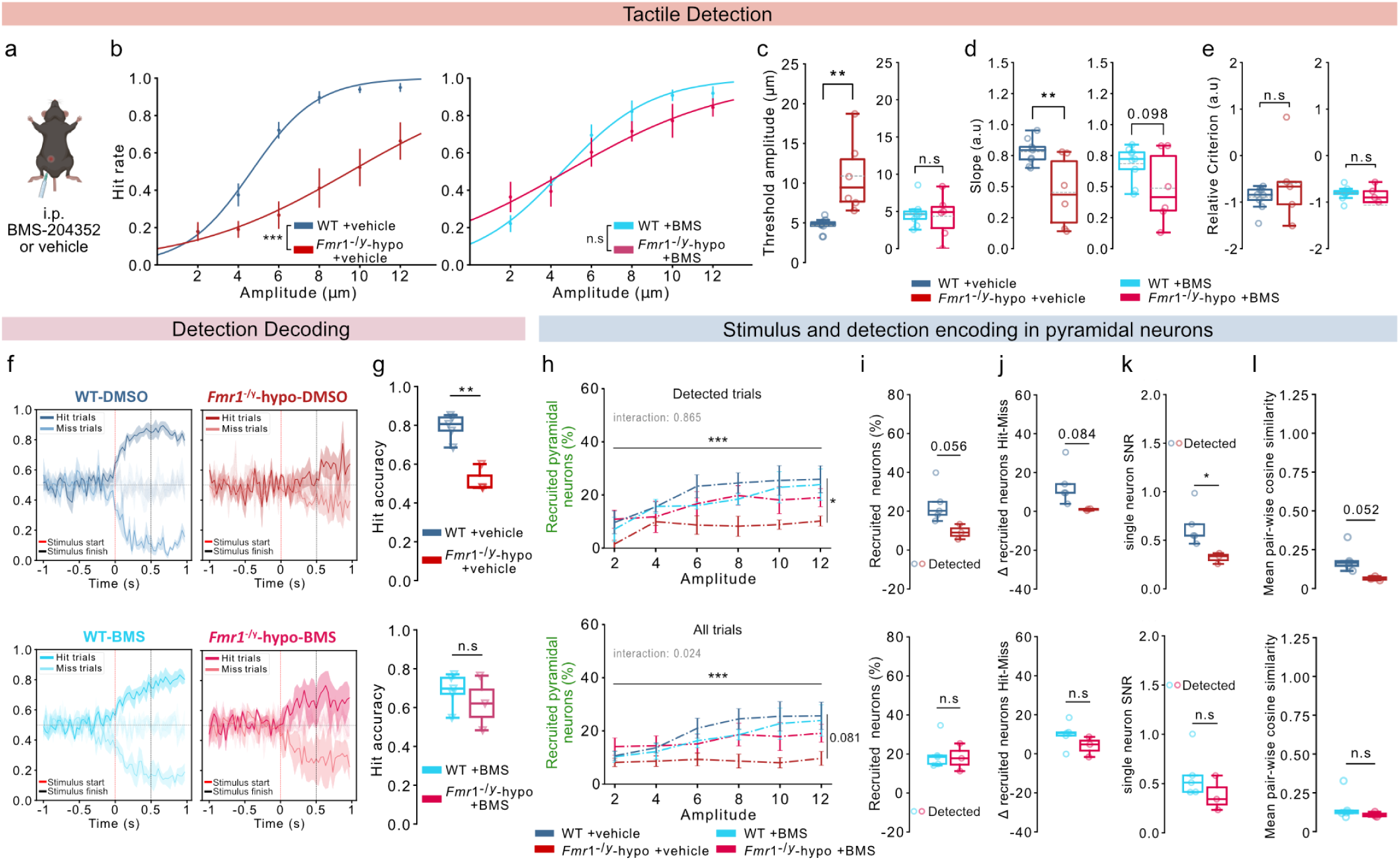
Correcting hyperexcitability improves tactile perception and detection decoding by enhancing stimulus and detection encoding in S1-FP of *Fmr1*^-/y^-hyposensitive mice. For panels **b,-e,**: n=9 WT, 6 *Fmr1*^-/y^–hyposensitive mice, ∼32 repetitions of each amplitude per mouse for vehicle, ∼37 repetitions for BMS. For panels **f,-l,**: n=5 WT, 3 *Fmr1*^-/y^–hyposensitive mice,1 session with ∼10 repetitions of each amplitude per mouse for vehicle and BMS. **a,** Intraperitoneal injection of BMS-204352 (BMS) or vehicle 30-45 minutes before the tactile detection task. **b,** Psychometric curves fitted on average Hit rates for each stimulus amplitude for sessions following vehicle injection (left) or BMS injection (right). Detection thresholds **(c,)** and perceptual accuracy (slope; **d,**) calculated based on the psychometric curves in **(b,)**. **e,** Relative criterion (strategy) for treatment with the vehicle (left) and BMS (right). **f,** Response decoding upon vehicle treatment in WT mice (top, left) and *Fmr1*^-/y^-hyposensitive mice (top, right) or upon BMS treatment in WT (bottom, left) and *Fmr1*^-/y^-hyposensitive mice (bottom, right). Stimulus start is indicated by a red dashed line and stimulus end by a black dashed line. The dark line represents true Hit classification and the light line represents false Miss classification. Classifiers were trained on each frame of prestimulus activity (1 s), tactile stimulation (500 ms), and post-stimulus activity (500 ms) to decode if the trial was detected (Hit) or non-detected (Miss). **g,** Comparison of the average Hit accuracy of the classifiers during stimulation between WT and *Fmr1*^-/y^-hyposensitive mice upon vehicle (top) or BMS (bottom) treatment. **h,** Average percentage of recruited pyramidal neurons for each stimulus amplitude in detected and non-detected trials (top) and detected-only trials (bottom). **i,** Proportion of recruited pyramidal neurons during detected trials upon vehicle (top) or BMS (bottom) treatment. Difference in the proportion of recruited pyramidal neurons **(j,)** and in the average z-score response **(k,)** between detected and non-detected trials upon vehicle (top) or BMS (bottom) treatment. **l,** Mean pair-wise cosine similarity of pyramidal trial-by-trial responses during stimulation upon vehicle (top) or BMS (bottom) treatment. P values were computed using Mixed ANOVA for panels **b, h,**; two-sided t-test for panel **c, d, e, g, i, j, k, l-top;** Mann-Whitney U test for panels **e, l-bottom,.** ***P<0.001, **P < 0.01, *P < 0.05 or n.s, not significant.

Pharmacological correction of hyperexcitability significantly improved tactile sensitivity in *Fmr1*^-/y^-hyposensitive mice (Fig. 7b), restoring their detection thresholds to levels comparable to those of their WT littermates (Fig. 7c). The BK_Ca_ channel agonist did not affect perceptual accuracy (psychometric slope; Fig. 7d) or the licking strategy of the *Fmr1*^-/y^-hyposensitive mice (Fig. 7e).

These results demonstrate that neuronal hyperexcitability in *Fmr1*^-/y^-hyposensitive mice plays a key role in their elevated detection thresholds and impaired tactile detection, and that targeted modulation of excitability can restore sensory perception.

### Neuronal hyperexcitability hinders the encoding and decoding of sensory detection in *Fmr1*^-/y^-hyposensitive mice

Next, we tested whether dampening hyperexcitability upon BMS treatment also improved the decoding reliability from S1-FP population activity. Using the same decoding classifiers as previously (Fig. 2), we examined the transformation of sensory input into perceptual output under vehicle and BMS conditions (Fig. 7f). Consistent with earlier results (Fig. 2d), vehicle-treated *Fmr1^-/y^*-hyposensitive mice exhibited reduced model accuracy for predicting detected trials compared to WT littermates (Fig. 7g-top). This impairment was abolished following BMS treatment, with decoding accuracy reaching WT levels (Fig. 7g-bottom), indicating improved neural representation of detection upon normalization of excitability.

To determine whether this improvement in decoding stemmed from enhanced detection encoding, we quantified the proportion of pyramidal neurons recruited during detected trials (Fig. 7h), as prior results showed a genotype effect for this cell type (Fig. S4d-e). Under vehicle treatment, *Fmr1^-/y^*-hyposensitive mice displayed a strong trend for reduced pyramidal recruitment (Fig. 7i-top), which was restored following BMS administration (Fig. 7i-bottom). This rescue was further supported by a normalized difference in neural recruitment between detected and non-detected trials in *Fmr1^-/y^*-hyposensitive mice after BMS treatment, comparable to that of WT controls (Fig. 7j). These results show that reducing hyperexcitability strengthens detection encoding via increased recruitment of pyramidal neurons, leading to better detection decoding.

Based on our finding of a strong correlation between neural recruitment and single-neuron SNR (Fig. 5e-f), we tested whether the BMS-induced increase in pyramidal neuron recruitment during detection was accompanied by an increased single-neuron SNR. Indeed, the diminished pyramidal single-neuron SNR observed under vehicle treatment (Fig. 7k-top) was ameliorated in *Fmr1^-/y^*-hyposensitive mice treated with BMS, showing no difference to that of WT mice (Fig. 7k-bottom). These findings demonstrate that correcting neural hyperexcitability in *Fmr1^-/y^*-hyposensitive mice enhances single-neuron SNR, thereby improving neural recruitment during detection.

Given that disrupted pyramidal ensembles contributed to impaired stimulus encoding in *Fmr1*^-/y^-hyposensitive mice (Fig. 4a), we next evaluated neural synchrony during stimulation. Vehicle-treated *Fmr1*^-/y^-hyposensitive mice showed a strong trend toward reduced pairwise correlation of pyramidal responses (Fig. 7l-top). This was significantly improved by BMS, which restored pyramidal neuron response correlations to WT levels (Fig. 7l-bottom), showing enhanced ensemble coordination during stimulus encoding.

Together, these findings demonstrate that reducing neuronal hyperexcitability in *Fmr1^-/y^*-hyposensitive mice increases single-neuron SNR, thereby enhancing both stimulus and detection encoding at the pyramidal population level. This enables reliable decoding of tactile perception from S1-FP population activity and restores tactile sensitivity.

## Discussion

In this study, we assessed how tactile detection and its neural basis are altered in autism by combining psychophysics with neuronal population imaging in a highly translational manner. We leveraged forepaw somatosensation–a system that is evolutionarily conserved across eutherian mammals including humans and non-human primates–to recapitulate the tactile symptoms of autistic individuals and directly link them to neocortical mechanisms.

We tackled the challenge of experimentally mimicking the nuanced and heterogeneous features of atypical sensory perception reported in autistic individuals in a transgenic mouse model, by reverse-translating a psychophysical task used in human studies^13,14,58^. Our results demonstrate a remarkable fidelity in capturing the nuances of altered sensory perception in autism, such as tactile hyposensitivity^13,14^, high interindividual variability^19,28–30^, and unreliable responses^8,39,40^. Consistent with human studies^31,59^, these tactile symptoms were not linked with cognitive deficits. Intriguingly, we reveal a subgroup of *Fmr1*^-/y^ mice with tactile hyposensitivity. This result was unexpected for an inbred mouse line but reinforces clinical reports of sensory subgroups in autism^31^ and underscores the importance of accounting for individual variability in experimental design and therapeutic targeting both in clinical and in preclinical studies. The elevated body weight of the hyposensitive *Fmr1*^-/y^ subgroup suggests that metabolic alterations in autism^36–38^ might impact the metabolic rate of the neocortex, thereby modulating cortical sensory gain and tactile sensitivity^34,35^.

Tactile perception alterations in autism depend both on the tactile sub-modality and the body part that receives the stimulus, revealing hypo- or hypersensitivity in autistic individuals depending on the context^13,14,16^. Our findings of tactile hyposensitivity in *Fmr1*^-/y^ mice are consistent with a previous study in this mouse model using a forepaw-dependent freely-moving discrimination task^60^. Importantly, tactile detection thresholds assessed through vibrotactile stimuli applied to the fingers are highly correlated with social difficulties^61^. Our results further underline the importance of the stimulated body part and the tactile sub-modality, as hindpaw or whisker stimulation has been shown to result in hypersensitivity^62,63^, increased affective reactivity^64^, and hyper-responsiveness^65^ in several mouse models of autism.

Our approach enabled a detailed investigation of the neural correlates underlying altered tactile perception in *Fmr1*^-/y^ mice. We show that neuronal responses in the forepaw-related primary somatosensory cortex (S1-FP) during stimulation were sufficient to predict behavioral responses in WT mice, congruent with results from the barrel cortex^66,67^ and primate studies^68^. Strikingly, this decoding was impaired in *Fmr1*^-/y^-hyposensitive mice. While multiple studies have assessed stimulus-evoked neural responses in autistic individuals and mouse models of the condition, we provide further insight into the encoding of both tactile stimuli and their detection in S1-FP, demonstrating for the first time that not only stimulus-evoked responses but also detection encoding is impaired in autism.

Reduced somatosensory responses have been previously reported in autistic individuals^18,19^ and whisker-related responses during passive stimulation were found decreased or degraded in mouse models of autism^69,70^. Delayed somatosensory processing has also been noted in some autistic individuals^71^, and reduced temporal precision in responses to whisker stimulation have been reported in young *Fmr1*^-/y^ mice^65^. Similarly, delayed auditory-evoked responses reflecting low-level encoding deficits have been described in individuals with autism and valproic acid (VPA)-exposed mice^72^. Delayed neuronal desynchronization in the somatosensory cortex of *Fmr1*^-/y^ mice during development is accompanied by network alterations that persist in adulthood^73^. Our findings demonstrate that in trained adult animals engaged in a perceptual task, stimulus-evoked neuronal correlations are reduced. These results suggest that early developmental hyper-synchrony may result in dysfunctional ensemble coordination in adulthood, potentially reflecting impaired circuit refinement. Consistently, reduced neural synchrony during auditory stimulation has also been documented in adolescents and adults with Fragile X Syndrome^74^.

Our findings extend this work by providing a mechanistic dissection of altered detection encoding and revealing three converging deficits in *Fmr1*^-/y^-hyposensitive mice: diminished recruitment of pyramidal neurons, disrupted temporal precision, and degraded ensemble coordination. Collectively, these deficits impair the population-level encoding of tactile stimuli and their detection in S1-FP. Whereas detection was not represented at the single-neuron level, reduced single-neuron SNR was tightly correlated with weakened population encoding—a critical determinant of sensory perception. This coupling identifies single-neuron SNR as a key mechanism linking cellular encoding to network-level stimulus representation and behavior.

We further linked these encoding deficits to neuronal hyperexcitability in S1-FP, previously reported in *Fmr1*^-/y^ mice^54,56,75^. Correcting neuronal hyperexcitability with a BK_Ca_ channel agonist^54–57^ restored tactile perception via improved stimulus and detection encoding. Hyperexcitability distorts canonical computations such as divisive normalization^76^, reducing the relative weight of stimulus-related input as it is normalized against elevated non-stimulus-related activity that rises from the hyperactive network. Such a hyperexcitable network contributes to decreased single-neuron SNR in *Fmr1*^-/y^-hyposensitive mice, which in turn diminishes the proportion of stimulus-recruited neurons. Moreover, hyperexcitability increases response variability, which has been proposed as a hallmark of autism^41–43,56,77^, evidenced in our data as less distinct responses between detected and non-detected trials and disrupted pyramidal ensemble dynamics. Treatment with a BK_Ca_ channel agonist reduces spontaneous activity and thus endogenous background noise^56^, thereby enhancing SNR and facilitating stimulus detection.

Together, our results identify the following mechanistic chain: neuronal hyperexcitability leads to reduced single-neuron SNR that in turn diminishes population-level stimulus and detection encoding and results in elevated detection thresholds and unreliable responses in *Fmr1*^-/y^-hyposensitive mice (Fig. 8).

**Figure 8.**
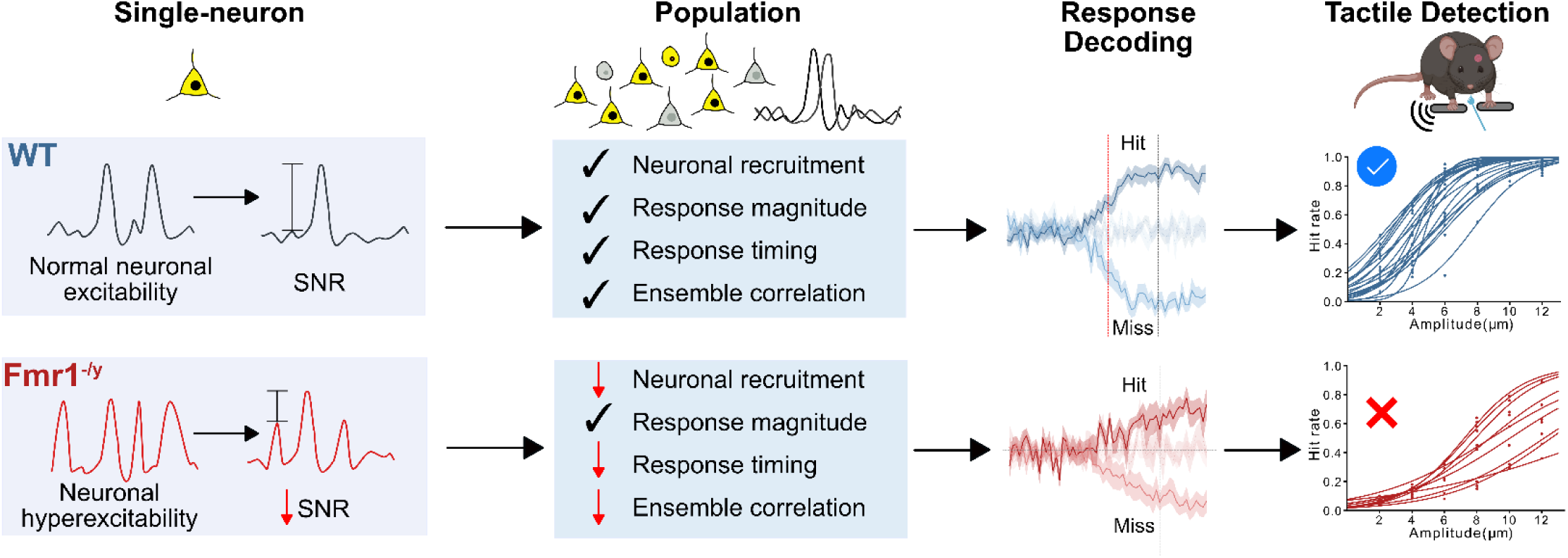
Schematic summary of altered tactile perception in *Fmr1*^-/y^-hyposensitive mice. Summary diagram illustrating the key behavioral, neuronal, and mechanistic findings. In WT mice, low-level tactile stimuli are reliably encoded in the forepaw primary somatosensory cortex (S1-FP) with high single-neuron signal-to-noise ratio (SNR), robust pyramidal neuron recruitment, precise response timing, and coordinated ensemble activity, enabling accurate response decoding and stimulus detection. In a tactile-hyposensitive subgroup of *Fmr1*^-/y^ mice, neuronal hyperexcitability leads to reduced single-neuron SNR that in turn drives diminished pyramidal recruitment, delayed responses, and weakened ensemble coordination, resulting in unreliable cortical decoding of detection and elevated behavioral thresholds.

In conclusion, we discover the neural mechanisms underlying neocortical encoding of low-level vibrotactile stimuli and their detection—highly relevant in the daily experience of humans and other mammals—and define how they change in autism. The translational value of our approach contributes to understanding how sensory stimuli and their detection are encoded in the brain and how perceptual decisions are formed. Our work further advocates for subject classification into subgroups based on sensory features, both in human studies^78–80^ and in preclinical models of autism. This stratification is crucial for evaluating novel treatments in preclinical and clinical studies. More broadly, our experimental platform enables the dissection of detection encoding across species, bridging physiology with perceptual decision-making. Moreover, it can be applied to other neurodevelopmental and psychiatric disorders where sensory symptoms are prevalent, including ADHD, sensory processing disorder, and schizophrenia^81^. By anchoring mechanistic studies to translationally relevant behaviors, this approach provides a direct path from circuit dysfunction to perceptual phenotype, paving the way for personalized interventions^82^ in sensory processing disorders.

## Materials and Methods

### Experimental design

We reverse-translated a task used in human studies^13,14^ and developed a novel Go/No-go detection task for vibrotactile stimuli which we combined with two-photon calcium imaging of excitatory and inhibitory neurons in the forepaw-related primary somatosensory cortex (FP-S1), to study altered tactile perception in autism and define its neural correlates.

### Ethical statement

All procedures adhered to EU directive 2010/63/EU and French law, approved by the Bordeaux Ethics Committee and Ministry for Higher Education and Research. Mice were housed in reversed 12 h light/dark cycle (light on at 21:00) under controlled conditions (22–24 °C, 40–60% humidity) with ad-libitum food and ad-libitum access to water before the water restriction period. Experiments were conducted during the dark phase under red light.

### Mice

Second-generation *Fmr1* knockout (*Fmr1*^−/y^)^27^ and wild-type littermate mice, 5-16 weeks old, were used. Mice were fully congenic on a C57Bl/6J background (backcrossed for >10 generations into C57Bl/6J). Male wild-type (*Fmr1*^+/y^) and knockout (*Fmr1*^−/y^) littermates were generated by crossing *Fmr1*^+/−^ females with *Fmr1*^+/y^ male mice from the same colony. Post-weaning, mice were group-housed (2–4 males per cage), balanced for genotype, and provided cotton nestlets and carton tubes. Genotypes were re-confirmed post hoc by tail-PCR.

### Surgery

Mice (P33-42) were anesthetized with isoflurane (4.5% induction, 1.5-2% maintenance), with anesthesia depth confirmed by absence of foot-pinch reflex and whisker movement. Mice were head-fixed using non-puncture ear bars and a nose clamp, and optic gel (Ocry-gel) was applied to both eyes. Body temperature was maintained at 37 °C with a heating pad and rectal probe. Pre-surgery analgesia consisted of Buprenorphine (s.c., 0.1 ml of 1:10 solution). Following hair trimming and Betadine antisepsis, local analgesia was induced (s.c., 0.1 ml of 1:4 Lidocaine-saline solution) and allowed to act for 2–5 min. The scalp was carefully removed and a 3 mm diameter craniotomy was made above the S1 forepaw region (0.75 mm anterior and -2.5 mm lateral from Bregma, confirmed with intrinsic imaging coupled with forepaw stimulation) using a dental drill (World Precision Instruments). The cortical surface was rinsed with saline throughout the surgery. Stereotactic injections of 100 nl of AAV1/2-syn-jGCaMP8m (titer: 2.6E+12 gcp/ml, diluted 1:4 in Hank’s Balanced Salt Solution) and AAV1/2-mDlx-HBB-chI-mRuby3-SV40p(A) (titer: 8.6E+13 gcp/ml, diluted 1:2 in Hank’s Balanced Salt Solution) were delivered (50 nl/min) in layers 2/3 (z depth 0.3mm). A double coverslip (3 mm lower glass glued to 4 mm upper glass with NOA 61 optical adhesive, Norland Products) was implanted over the craniotomy and sealed with cyanoacrylate glue. A head-post was attached with cyanoacrylate and dental cement. OptiBond Universal (Kerr) and Charisma dental filling material (Kulzer) were applied and cured with LED blue light, followed by a dental cement cap. Mice recovered on a warmed blanket for 1 hour post-anesthesia.

### Go/No-Go vibrotactile decision-making task

#### Setup

The vibrotactile decision-making setup was positioned in an isolation cubicle to minimize sound and light interference during the experiment. Mice were placed in a body tube and were head-fixed with their forepaws resting on two steel bars (6 mm diameter, Thorlabs). The right bar was mounted to a Preloaded Piezo Actuator equipped with a strain gauge feedback sensor and controlled (P-841.6 and E-501, Physik Instrumente) in a closed loop, as described before^83^. Water reward was delivered through a metal feeding needle (20G, 1,9mm tip, Agntho’s AB) connected to a lickport interface with a solenoid valve (Sanworks) equipped with a capacitive sensor (https://github.com/poulet-lab/Bpod_CapacitivePortInterface). The behavioral setup was controlled by Bpod (Sanworks) through scripts in Python (PyBpod, https://pybpod.readthedocs.io/en/latest/).

#### Habituation to head-fixation and water restriction

Mice (P40-P50) were handled using carton tubes and the cupping technique until they became comfortable with the experimenter. Mice were gradually habituated to the experimental setup and head fixation for 5 days. On day 3, a water-restriction protocol was implemented, providing access to liquid water during sessions and a solid water supplement (Hydrogel, BioServices) in their home cages. Animals received 1.5–2 ml water/day (50-65% of ad-libitum intake), without exceeding 10% body weight loss. Each mouse had access to 6-8 g of Hydrogel on weekends (100% of ad-libitum intake). Water restriction continued through training and testing.

#### Go/No-Go vibrotactile task training and testing

Habituated 8-week-old mice were trained to associate a vibrotactile stimulus (pure sinusoid, 500 ms duration, 15 µm amplitude, 40 Hz frequency) with a water reward (8 µl).

Go trials consisted of stimulus delivery followed by a 2-s response window during which the mice could lick to receive the reward. No-Go (catch) trials had no stimulus, and licking triggered a 5-s timeout. Inter-trial intervals varied between 5-10 s.

Training included three phases:

a. Automatic water delivery at the response window start.
b. Pre-training: lick-triggered water delivery on Go trials and timeout on No-Go trials.
c. Training: lick-triggered reward only if mice refrained from licking during the 3–8 s inter-trial interval before stimulus delivery; licking during this interval or on No-Go trials resulted in a 5 s timeout.

Sessions consisted of 300 pseudorandomized trials: 70–80% Go, 20–30% No-Go. Pilot experiments with an extra sensor to monitor forepaw placement confirmed that the mice did not remove their forepaws from the bar before stimulus delivery.

Pre-training criteria: ≥80% successful Go trials, <40% spontaneous licking during No-Go. Training criteria: ≥80% successful Go and <30% unsuccessful No-Go trials averaged over 3 days.

Mice meeting criteria were tested on novel vibrotactile amplitudes (2–12 µm, 10 Hz, 500 ms, 6 stimuli per session) with a 90% Go:10% No-Go ratio, presented pseudorandomly.

#### Go/No-Go vibrotactile task analysis

Behavioral data were analyzed using custom Python scripts. Four main metrics were calculated from lick events:

a. Hit rate: Hits ÷ total Go trials (stimulus delivered)
b. Miss rate: Misses ÷ total Go trials (stimulus delivered)
c. Correct Rejection rate: Correct rejections ÷ total No-Go trials
d. False Alarm rate: False alarms ÷ total No-Go trials

Prestimulus spontaneous licking was quantified as the number of timeouts during Go trials divided by total Go trials. Training duration was the total days to pass pre-training and training phases.

Only testing sessions with <40% unsuccessful No-Go trials were included. Psychometric curves were fitted on the Hit rate for each stimulus amplitude using a general linear model, averaging ∼140 repetitions per amplitude. Detection thresholds were derived as the stimulus amplitude at the sigmoid inflection point. Accuracy was assessed by the psychometric curve slope. Relative criterion was calculated as:

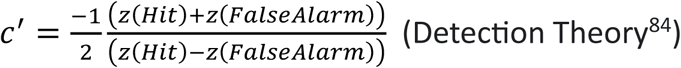

where z(Hit) is the standard deviation unit for the Hit rate and z(False Alarm) is the standard deviation unit for the False Alarm rate.

#### Trial-by-trial variability

Behavioral variability was quantified as the variance of Hit responses across Go trials for each stimulus amplitude. Low variance indicates consistent detection or non-detection of a stimulus. Variability was normalized and calculated at each mouse’s threshold amplitude (x), as well as the immediately suprathreshold (x + 2 µm) and subthreshold (x – 2 µm) amplitudes.

#### Clustering of *Fmr1^−/y^* mice in subgroups

The sci-kit learn implementation of k-means was used on the Hit rates of *Fmr1^−/y^* mice across all stimulus amplitudes, with 2 clusters targeted and the default parameters.

### Two-photon calcium imaging

#### Setup

Calcium measurements were performed using a Femtonics FEMTOSmart microscope coupled with a widely tunable, femtosecond Ti:Sapphire Laser (Chameleon Ultra, Coherent) pumped by a solid-state laser (Verdi 18W, Coherent). Excitation at 920 nm was used for GCaMP8m, and at 1000 nm for mRuby3, and two photomultiplier tubes were used for the collection of green and red fluorescence, respectively. Laser power was modulated using a Pockels cell A 20x/1 NA objective (Zeiss) was used to obtain 512 × 512-pixel images covering 480 × 480 μm of cortex. Images were acquired at 30.96 Hz using the resonant scanner system (Femtonics). Measurements were written as a series of 32-bit images and saved as ome files. Analog and digital inputs sent by Bpod and the piezoelectric sensor controller were collected and saved as timestamps by the microscope and were used to synchronize the behavioral events with the calcium traces from imaging.

#### Analysis

Imaging data were acquired as 2000-frame ome files and converted to TIFF using Fiji (ImageJ). Motion correction, region of interest (ROI) detection, and fluorescence extraction were performed using Suite2p (https://suite2p.readthedocs.io/en/latest/index.html) with standard parameters, double registration, and denoising. Neuron selection was manually curated based on ROI shape and calcium traces. Inhibitory neurons were identified by colocalizing green and red fluorescence channels in ImageJ. Subsequent analyses were performed separately for excitatory and inhibitory neurons. Custom Python scripts synchronized behavioral and imaging data and calculated ΔF/F and z-scores. Neuropil contamination was corrected in all traces prior to ΔF/F and z-score calculations as:

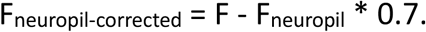

where F is the fluorescence of the neuron (ROI) and F_neuropil_ is the fluorescence of the neuropil associated with the neurons, as calculated by suite2p.

To calculate ΔF/F, baseline fluorescence (F) was calculated for each neuron as the average fluorescence during the 10 s period with the lowest variation (STD) in F.

z-scores were calculated as:

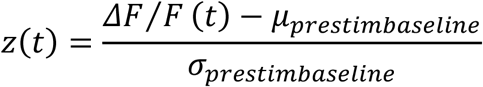

where *µ_prestim baseline_* is the mean ΔF/F during the 1 s before stimulation across all trials and σ_prestim baseline_ is the standard deviation ΔF/F during 1 second before stimulus

All stimulus-specific response analyses were restricted to the stimulus period before the animal’s first lick, preventing contamination by movement-related activity.

### Stimulus-related responses

For each trial, a neuron’s stimulus-related activation was assessed by comparing the 85th percentile of its z-score during stimulation to 1,999 randomly selected time points from the recording (excluding stimulus and reward periods). If the z-score during stimulus delivery exceeded the 95th percentile of these random points, the neuron was classified as stimulus-activated.

Stimulus-related inhibition was determined similarly, using the 10th percentile of the stimulation z-score compared to the 95th percentile of the random points, with neurons considered inhibited if their z-score was lower than -1 times this value.

This randomization test follows Prsa et al^83^ and uses a significance threshold of P < 0.05.

### Response characterization (peak amplitude, peak delay)

Response amplitude was calculated for all neurons that were activated or inhibited during stimulus delivery as the maximum or minimum z-score value during the stimulation window before the first lick. The peak delay was defined as the number of frames between the beginning of stimulus delivery and the peak response amplitude.

### Ensemble dynamics - Correlation of neuronal responses

To estimate the correlation of neuronal response patterns, neurons were represented as vectors of trials, with each trial assigned a value reflecting the neuron’s response to the stimulus: activated (1), inhibited (-1), or non-responsive (0). Cosine similarity was computed between each pair of neurons. For group comparisons, the pairwise similarity matrices were averaged (excluding the diagonal) to obtain a single cosine similarity value for each animal.

### Excitation-inhibition (E/I) ratio

E/I ratio was calculated for each trial as the number of activated pyramidal neurons divided by the number of activated GABAergic interneurons.

### Signal-to-Noise Ratio (SNR)

The single-cell SNR was computed by calculating the absolute difference between the neuron’s mean activity during each trial (z-scored) and its mean activity during catch trials. The population SNR was computed as the ratio of the percentage of recruited neurons during each trial to the mean percentage of recruited neurons during No-Go (catch) trials.

### Decoding of behavioral responses

Dynamic decoding classifiers were adapted from ref.^44^, using a logistic regression (Scikit-learn implementation) model with an L2 penalty term to the. The regularization strength (C) was determined through double cross-validation, choosing from values of 0.0001, 0.001, 0.01, 0.1, and 1, as described in ref.^85^. Classification accuracy was computed per frame, with shaded areas showing the SEM. Decoding accuracy for time windows was calculated by averaging frame accuracies for each time window across trials. To balance detected/undetected trials (due to altered Hit/Miss ratios in WT and *Fmr1*^−/y^ mice), random undersampling was applied (Scikit-learn), retaining random trials from the majority class to match the minority class size.

Each mouse’s data trained a model to classify trial outcomes (Hit vs. Miss). Four-fold cross-validation was performed per session, training a new model for each fold and time point; reported accuracy is the test average across folds. A separate model was trained for each imaging frame, using single a scalar ΔF/F value for each cell of all trials on a given frame as the input vector.

To assess the model’s performance, shuffled data were used for comparison. These shuffled datasets were created by randomly reassigning the labels of the training data, allowing for an estimate of the model’s performance under the null hypothesis of no relationship between the input and the outcome.

To compare decoding accuracy between neuron types, separate models were trained for excitatory (EXC) and inhibitory (INH) neurons, and their accuracies, obtained using 4-fold cross-validation, were compared to that of the full dataset.

### Pharmacology

For rescue experiments, vehicle (0.9 M NaCl supplemented with 1.25% DMSO and 1.25% Tween 80) or BMS-204352 (2 mg/kg, Tocris) was administered via intraperitoneal injection 30-45 minutes before behavioral testing (as described previously^54^). All mice were first habituated to restriction. Each mouse received 2 days of vehicle followed by 2 days of BMS-204352 treatment. Go/No-Go vibrotactile task testing occurred 30-45 minutes after each injection. All sessions were included in the analysis of behavioral results, one vehicle and one BMS session were chosen for the analysis of neuronal activity measurements.

### Histology

Following the end of the experiments, mice were anesthetized with isoflurane (4.5%) and euthanized by cervical dislocation. Brains were extracted, fixed in 4% PFA overnight, and washed in 0.1 M phosphate buffer. Coronal slices of 100 mm thickness were obtained using a vibrotome (Leica VT1000 S). Slices were stained with DAPI, and viral injection sites in the forepaw primary somatosensory cortex were confirmed via fluorescence microscopy (Nikon Eclipse Ti-U).

### Statistics

Data are presented as mean ± s.e.m. Box plots show the median, interquartile, range, mean and individual values. Sample sizes are indicated in figure legends. Statistical analyses were performed in Python using SciPy and Pingouin. Normality was assessed for all datasets. For comparisons between two groups, two-tailed paired or unpaired t-tests were used when normality was confirmed; otherwise, Mann-Whitney U tests (between groups) or Wilcoxon Signed Rank tests (within groups) were applied. When more than two groups were present, effects were evaluated with one-way ANOVA or Kruskal-Wallis ANOVA. Genotype and Amplitude or Variance effects were tested with Mixed ANOVA, or the Friedman test was used for repeated measures analysis. For the percentage of recruited neurons over amplitude, the data were transformed using the Yeo-Johnson transformation before performing the Mixed ANOVA to meet the normality assumption. Normality and homoscedasticity were assessed using QQ plots and Scale-Location plots.

### Blinding and randomization

All experiments and analyses were conducted in a blinded manner. Although formal randomization was not applied, sample selection was effectively random due to blinding. For pharmacological rescue experiments, equal numbers of *Fmr1^−^*^/y^ and wild-type mice received BMS-204352 or vehicle. Treatments were evenly assigned within littermate groups.

### Code availability

Custom-made Python codes used in this study can be found on the following GitHub repository: https://github.com/OuraniaSem/Percephone.git

### Data availability

The raw data generated in this study have been deposited on the figshare database: https://doi.org/10.6084/m9.figshare.29900456

Source data are provided with this paper.

## Acknowledgments

We would like to thank the animal facility and the genotyping platform of the NeuroCentre Magendie (INSERM U1215 Unit) for assisting in animal breeding, maintenance, and genotyping. We thank Apolline Joliot for assisting in immunohistochemistry experiments. We thank Dr. James Poulet (Max-Delbrück-Centrum für Molekulare Medizin, Berlin) for providing the capacitive sensors for the behavioral task. We thank Dr. Naoya Takahashi (Interdisciplinary Institute for Neuroscience, Bordeaux) and Dr. James Poulet (Max-Delbrück-Centrum für Molekulare Medizin, Berlin) for providing feedback on a previous version of the manuscript. Mouse schemas were made using Biorender. This project was funded by ANR-MultiSens (A.F.), Foundation for Medical Research Postdoc Fellowship (O.S.), INSERM (A.F.), CNRS (YLF), ANR-EMT-Sens (YLF), Marcel Dassault-Fondation FondaMental Award 2019 (A.F.), Simons Foundation Autism Research Initiative (A.F.), GIS Autisme & TND (O.S.), Fondation FondaMental (A.F.), 2024 NARSAD Young Investigator grant - Grant ID: 32694 (O.S.).

## Author Information

### Contributions

A.F. and M.G. conceived the project. A.F., M.G., and O.S. designed the experiments. O.S. performed the experiments. A.S.J. and Y.L.F. contributed to tactile responsivity experiments and their analysis. A.C. wrote the Python code for the behavioral experiments. O.S., T.G., and C.V. analyzed the data and wrote the necessary Python scripts. O.S. prepared the figures. O.S., T.G., and A.F. interpreted the data. O.S. wrote the manuscript, A.F., T.G., and C.V. provided feedback, and A.F. refined the manuscript. M.G. and Y.L.F. provided feedback on a previous version of the paper.

## Ethics declarations

### Inclusion and diversity statement

We support inclusive, diverse, and equitable conduct of research. Throughout the text, tried to use inclusive language as much as possible and terms preferred in the autistic community and are less stigmatizing.

