## Supplemental Data for "Diminished signal-to-noise ratio disrupts somatosensory population encoding and drives tactile hyposensitivity in the *Fmr1*^-/y^ autism model"

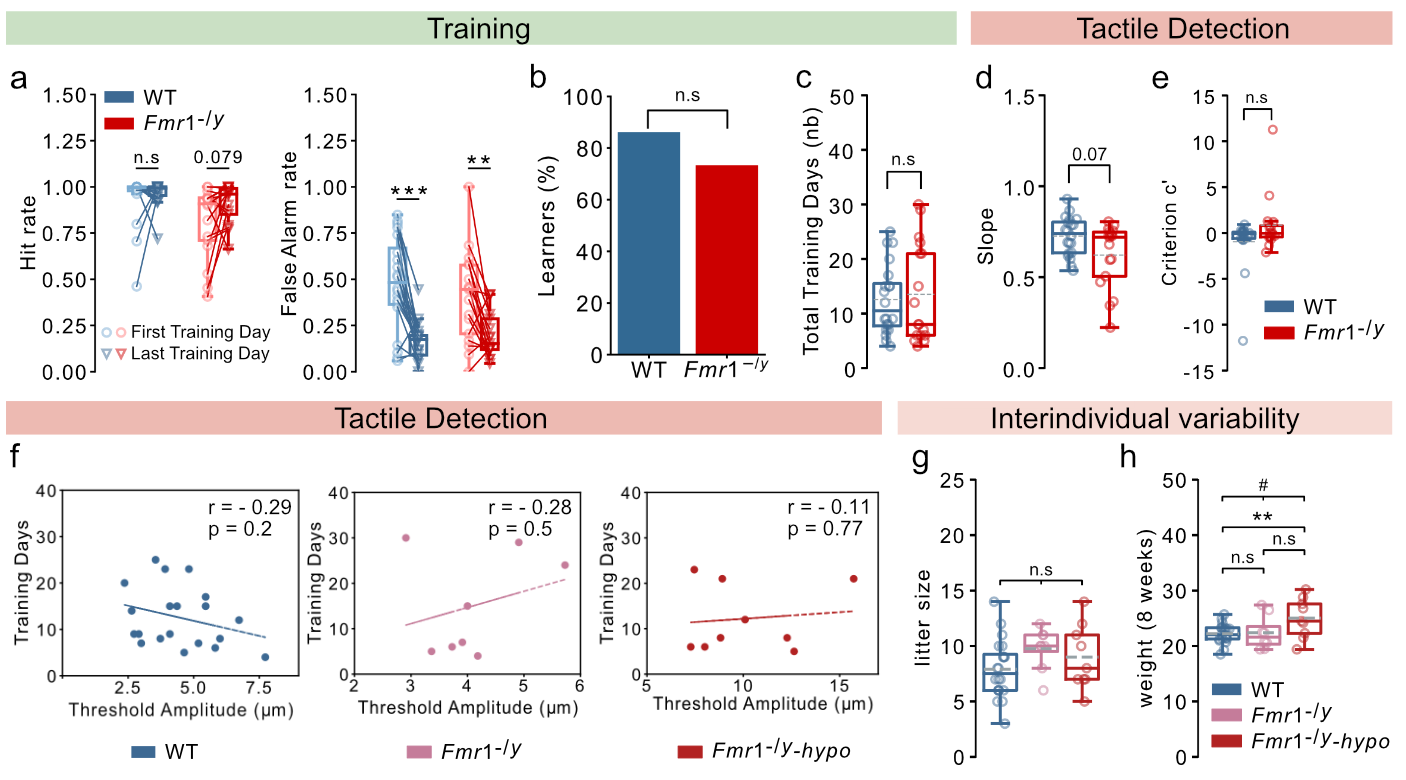

**Figure S1.** **Learning, tactile detection, and subgroup features**

For panels **a, c, d, f, g, h,**: n=20 WT, 17 *Fmr1*^-/y^ mice (of which 9 *Fmr1*^-/y^-hyposensitive). For panel **b**,: n=29 WT, 30 *Fmr1*^-/y^ mice. For panel **e,**: n=19 WT, 16 *Fmr1*^-/y^ mice. **a,** Hit rate (left) and False Alarm rate (right) during the first day of training (following pre-training, see Methods; light-color, circles) and the last day of training (dark color, triangles) for all mice that underwent detection testing. **b,** Percentage of mice that reached the learning criterion. **c,** Total number of days spent in pre-training and training for all the animals that underwent detection testing. **d,** Perceptual accuracy calculated as the slope of the psychometric curve for each mouse. **e,** Licking strategy calculated as relative criterion c’ for each mouse. **f,** Correlation of the total number of pre-training and training days with the threshold amplitude for WT (left), typically-detecting *Fmr1*^-/y^ mice (middle), and *Fmr1*^-/y^-hyposensitive mice (right). **g,** Size of the litter each mouse was born in. **h,** Weight of each mouse at 8 weeks before water restriction. P values were computed using a Wilcoxon signed-rank test and two-sided paired t-test for panel **a,;** a Chi square for panel **b,**; Mann-Whitney U test for panels **c,** **d,** **e,;** Pearson correlation coefficient for panel **f,;** and an one-way ANOVA and two-sided t-test for panels **g, h**,; two-sided paired T-test for panel **a,**; and two-sided T-test for panels **g, h,**. ***P < 0.001, **P < 0.01 ^#^P < 0.05, or n.s, not significant. ^#^ indicates ANOVA results.

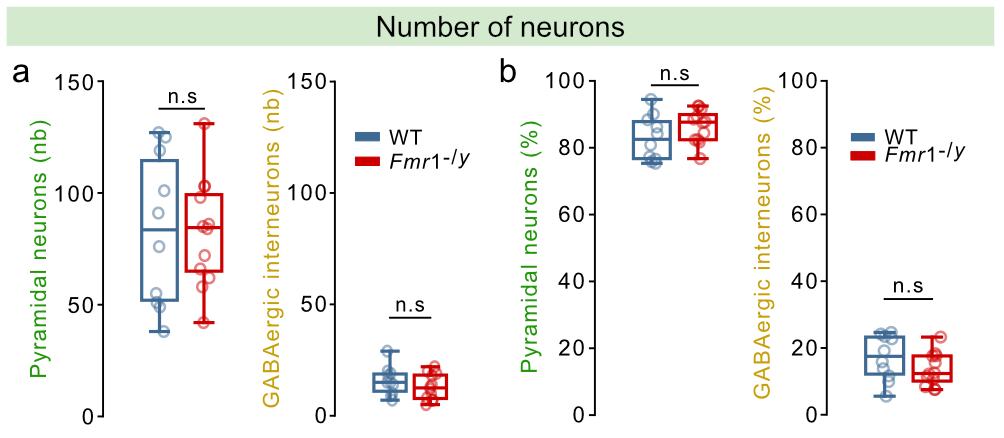

**Figure S2.** **Number of recorded neurons.** For all panels n=10 WT, 12 *Fmr1*^-/y^ mice. **a,** Number of pyramidal neurons (left) and GABAergic interneurons (right) detected in the field of view as regions of interest (ROIs) after manual curation. **b,** Same as **a,** but expressed as percentage of the neurons in the field of view. P values were computed using two-sided t-test for all panels. n.s, not significant.

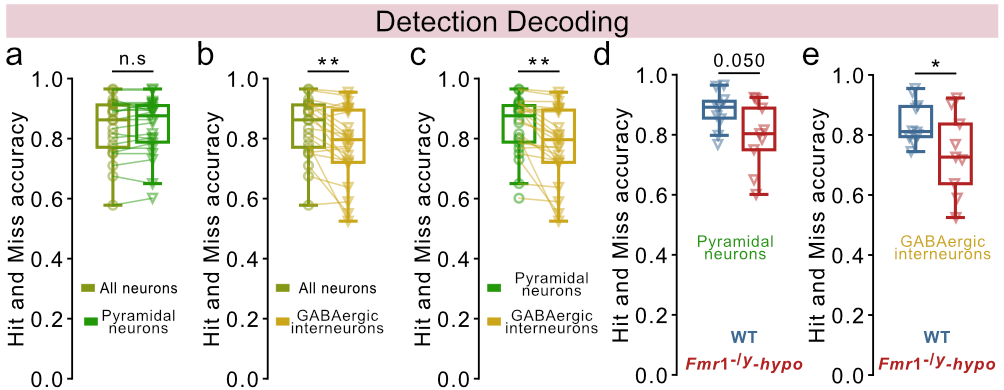

**Figure S3. Detection decoding in S1-FP based on pyramidal neurons or GABAergic interneurons.** For panels **a-c**: 19 mice (10 WT, 9 *Fmr1*^-/y^–hyposensitive mice). For panels **d, e,**: n=10 WT, 9 *Fmr1*^-/y^–hyposensitive mice. ~10 repetitions of each amplitude per mouse. Classifiers were trained on the mean neuronal activity during stimulation (500 ms) to decode if the trial was detected (Hit) or non-detected (Miss). **a,** Comparison of decoding accuracy for Hit and Miss trials when training the model with all neurons versus only pyramidal neurons. **b**, Same as **a,** but comparing the model’s accuracy when trained with all neurons versus only with GABAergic interneurons. **c**, Same as **a,** but comparing the model’s accuracy when trained with pyramidal neurons versus with GABAergic interneurons. **d**, Decoding accuracy for Hit and Miss trials when training the model with pyramidal neurons in WT and *Fmr1*^-/y^-hyposensitive mice. **e**, Same as **d**, but using only GABAergic interneurons. P values were computed using two-sided paired t-test for panels **a, b, c,**; and two-sided t-test for panels **d, e,**. **P < 0.01, *P < 0.05, or n.s, not significant.

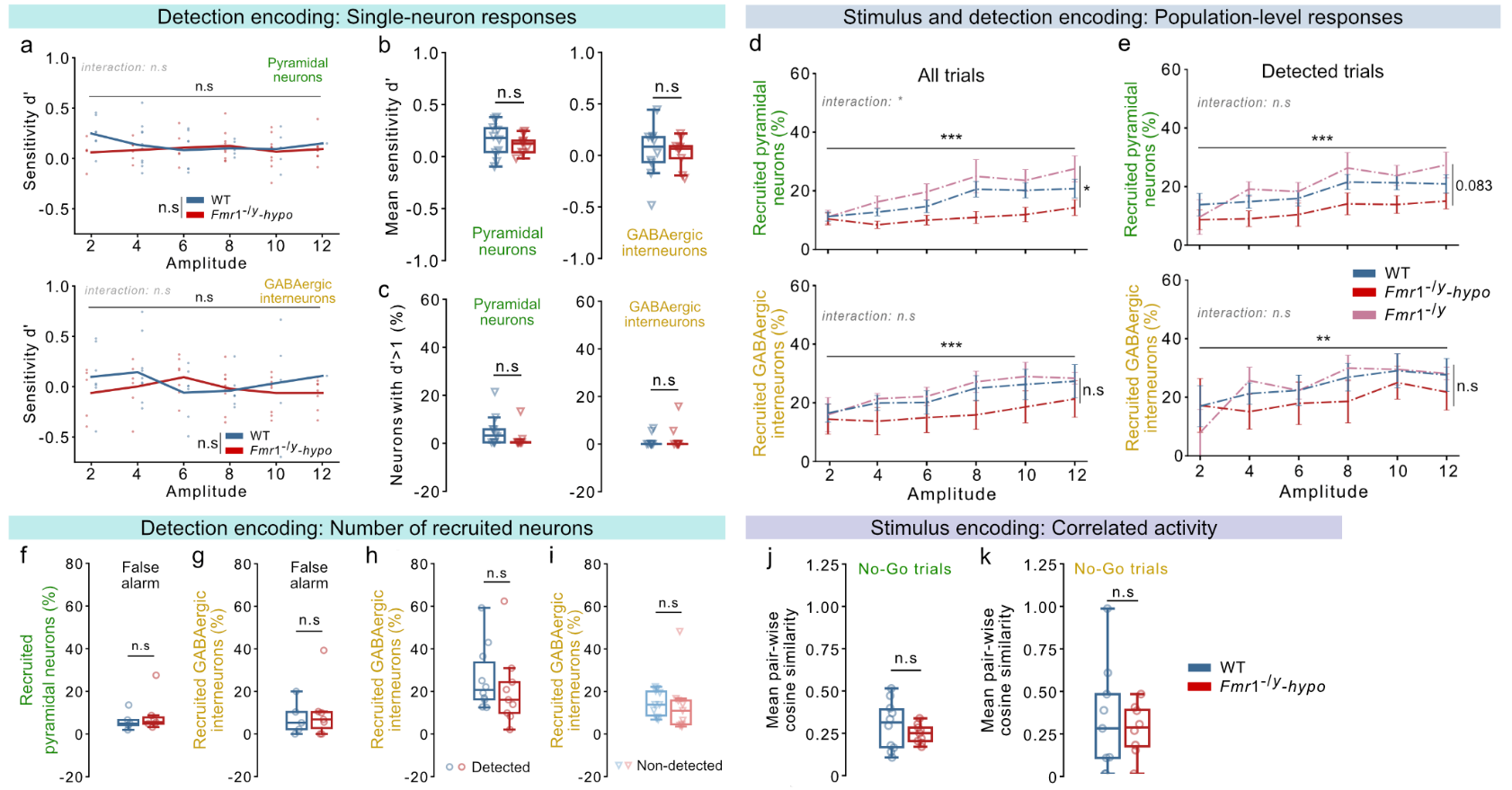

**Figure S4. Stimulus and detection encoding in S1-FP.**

For panels **a**: n=1-10 WT, 5-9 *Fmr1*^-/y^-hyposensitive mice, depending on how many mice had detected and non-detected trials at each stimulus amplitude. For panels **b, c, h, i, j**: n=10 WT, 9 *Fmr1*^-/y^-hyposensitive mice. For panels **d-e**: n=10 WT, 3 *Fmr1*^-/y^ mice with intact detection thresholds, 9 *Fmr1*^-/y^-hyposensitive mice. For panels **f, g,**: n=5 WT, 7 *Fmr1*^-/y^-hyposensitive mice. For panel **k,**: n=9 WT, 8 *Fmr1*^-/y^-hyposensitive mice. 1 session of ~10 repetitions of each amplitude per mouse. **a,** Mean single-neuron detection sensitivity (d’) for stimulus-recruited pyramidal neurons (top) and GABAergic interneurons (bottom) at each stimulus amplitude. **b,** Same as **a,** but for with all stimulus amplitudes grouped together. **c,** Proportion of pyramidal neurons (left) and GABAergic interneurons (right) with detection sensitivity d’>1. **d,** Proportion of pyramidal neurons (top) and GABAergic interneurons (bottom) recruited (activated or inhibited) during stimulus delivery at each different amplitude. **e,** Same as **d,** but only for detected trials of each amplitude. **f,** Proportion of recruited pyramidal neurons during False Alarm No-Go trials. **g,** Same as **f,** but for GABAergic interneurons. **h,** Proportion of recruited GABAergic interneurons during detected stimuli. **i,** Same as **h,** but for non-detected stimuli. **j,** Mean pair-wise cosine similarity of the trial-by-trial responses of pyramidal neurons during No-Go (catch) trials. **k,** Mean pair-wise cosine similarity of the trial-by-trial responses of GABAergic interneurons during No-Go (catch) trials. P values were computed using a Mixed ANOVA for panel **a,**; a Mixed ANOVA after a Yeo-Johnson transformation of the data for panels **d, e,**; two-sided t-test for panels **b, h, j, k,**; Mann-Whitney U test for panels **c, f, g, i,**. ***P < 0.001, **P < 0.01, or n.s, not significant.

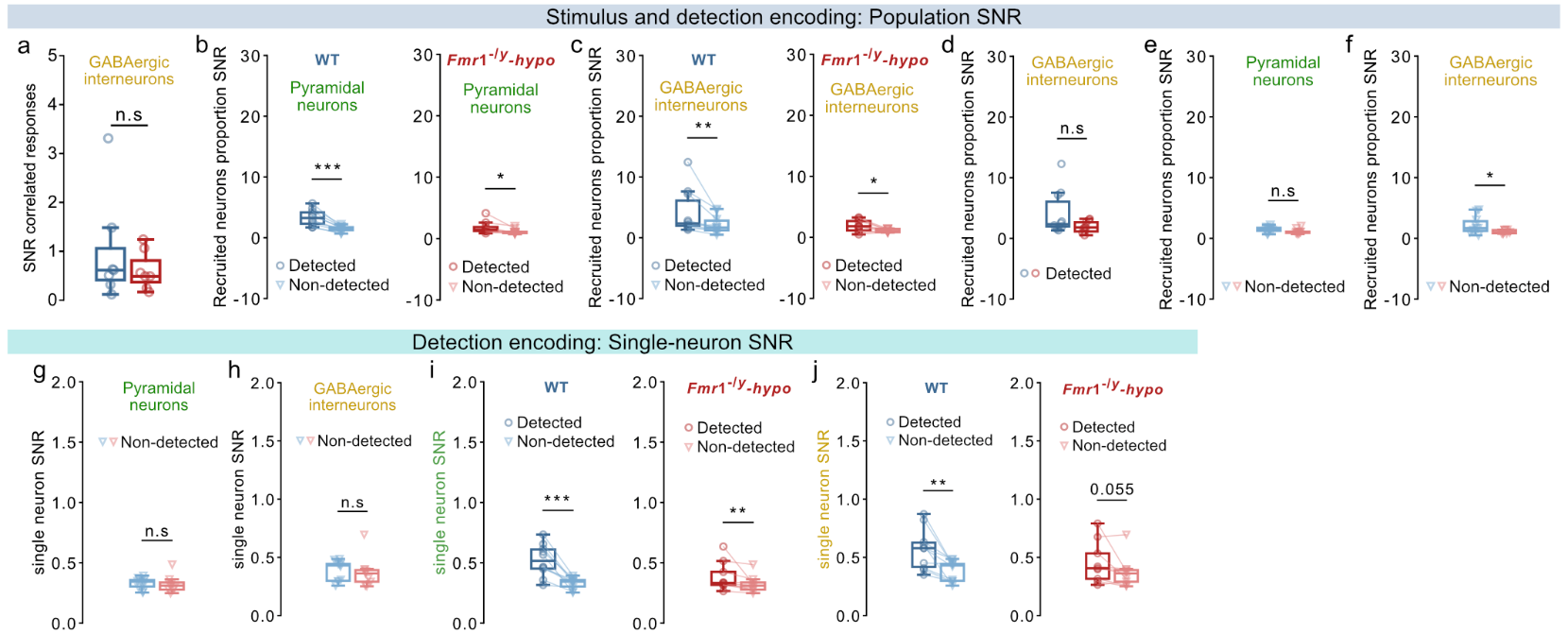

**Figure S5. Signal-to-noise ratio (SNR) during stimulus and detection encoding.**

For panel **a,**: n=7 WT, 7 *Fmr1*^-/y^-hyposensitive mice. For all other panels: n=10 WT, 9 *Fmr1*^-/y^-hyposensitive mice; 1 session of ~10 repetitions of each amplitude per mouse. **a,** Ratio of the mean pair-wise correlation of GABAergic interneurons during the tactile stimulus and during No-Go (catch) trials. **b,** Comparison of the ratio of recruited pyramidal neurons during detected trials to No-Go (catch) trials versus the ratio during non-detected trials to No-Go trials in WT (left) and *Fmr1*^-/y^-hyposensitive mice (right). **c,** Same as **b,** but for recruited GABAergic interneurons. **d,** Ratio of the proportion of recruited GABAergic interneurons during detected trials and during No-Go (catch) trials. **e,** Ratio of the proportion of recruited pyramidal neurons during non-detected trials and during No-Go (catch) trials. **f,** Same as **e,** but for GABAergic interneurons. **g,** Difference between the average z-score response of pyramidal neurons during non-detected trials and during No-Go (catch) trials. **h,** Same as **g,** but for GABAergic interneurons. **i,** Comparison of the difference between the average z-score response of pyramidal neurons during detected trials to No-Go (catch) trials versus the difference during non-detected trials to No-Go trials in WT (left) and *Fmr1*^-/y^-hyposensitive mice (right). **j,** Same as **i,** but for GABAergic interneurons. P values were computed using two-sided t-test for panels **f, g,**; a Wilcoxon signed-rank test for panels **b-right, c-left, i-right, j-right,**; two-sided paired t-test for panels **b-left, c-right, i-left, j-left ,**; Mann-Whitney U test for panels **a, d, e, h,**. ***P < 0.001, **P < 0.01, *P < 0.05, or n.s, not significant.
